# Redox-dependent extracellular interaction networks of Cysteine-rich Receptor-Like Kinases

**DOI:** 10.64898/2026.03.26.714216

**Authors:** Sergio Martin-Ramirez, Ran Lu, Mark Roosjen, Jente Stouthamer, Sjef Boeren, Daphne Homsma, Alejandro Thérèse Navarro, Jan Willem Borst, Judith Lanooij, Jan Maika, Rüdiger Simon, Willem Vermijs, Corine Geertsema, Monique van Oers, Adam G. Mott, Elwira Smakowska-Luzan

**Affiliations:** Laboratory of Biochemistry, Wageningen University & Research, Wageningen, The Netherlands; Biostatistics department, Rijk Zwaan, De Lier, The Netherlands; Plant Breeding Research Group, Wageningen University & Research, Wageningen, The Netherlands; Institute for Developmental Genetics, Heinrich Heine University, Düsseldorf, Germany; Laboratory of Virology, Wageningen University & Research, Wageningen, The Netherlands; Department of Cell & Systems Biology, University of Toronto, Toronto, Canada

## Abstract

Reactive oxygen species (ROS) regulate plant development and immunity, but how extracellular ROS signals are decoded and whether the Cysteine-rich Receptor-like Kinases (CRKs) truly serve as the long-suspected ROS sensors remains uncertain. Here, we combine high-throughput interactomics, redox proteomics, structural modelling, and genetics to map a ROS-dependent CRK interaction landscape in *Arabidopsis thaliana*. Using a redox-dependent interactome assay (RIA^CRK^) on 40 CRK extracellular domains (ECDs), we identified ROS-modulated dimerisation networks with enhanced inter-community connectivity and hub redistribution in the presence of ROS. Integrating this with developmental and flg22-induced expression profiles reveals spatiotemporally limited subnetworks that likely function during activated immunity and leaf senescence, both of which are associated with extensive ROS production. Differential cysteine alkylation coupled with mass spectrometry shows that cysteines in a subset of CRK ectodomains undergo ROS-dependent oxidation. Notably, solvent-exposed, vicinal cysteines C228/C229 in CRK28 emerge as prime redox-sensitive candidates. CRK28 homodimerises and heterodimerises with CRK17 in vivo, and mutation of C228/C229 retains plasma membrane localisation but abolishes CRK28 homodimerisation, indicating a redox-controlled dimerisation switch. Loss of CRK28 delays senescence, while CRK28 overaccumulation from its native promoter causes dwarfism, premature senescence, autoimmune-like phenotypes, extensive phosphoproteome rewiring, and associations with Pathogenesis Related (PR) proteins, ROS-detoxifying enzymes, receptor(-like) kinases, and vesicle trafficking components. These results indicate that CRK28 is a potential ROS-regulated hub connecting extracellular redox signals to CRK network organisation, immune response, and age-related senescence.

## Introduction

Throughout their lifespan, plants must precisely and appropriately perceive a range of overlapping external stimuli to ensure proper growth and development. To do this, plants developed multiple strategies through which signalling molecules are perceived by sensing proteins to initiate orchestrated responses. The generation of Reactive Oxygen Species (ROS) is one of the most common responses that plants exhibit in response to environmental challenges and as part of their developmental program ^1^. ROS are produced intra- and extracellularly, and their accumulation and localization is precisely controlled ^2,3^. ROS include singlet oxygen (^1^O_2_), superoxide (O_2_^•−^), hydroxyl radicals (OH^•^), and hydrogen peroxide (H_2_O_2_), the latter being the most stable. Basal ROS levels are essential for various cellular processes, including cell proliferation, differentiation, development, stress adaptation, and defence responses, among others ^4–12^. The production of ROS occurs in various subcellular compartments, including chloroplasts, mitochondria, peroxisomes, and the apoplast (extracellular space). The main processes responsible for producing intracellular ROS include mitochondrial respiration, chloroplastic photosynthesis, and peroxisomal photorespiratory reactions ^13–16^.

Apoplastic ROS accumulation is often caused by membrane-localized NADPH oxidases, such as the respiratory burst oxidase homologs (RBOHs) and other oxidases, activated by Receptor-Like Kinases (RLKs) during diverse cellular processes ^17–19^. RLKs canonically perceive various endogenous and exogenous molecules in the apoplast via their extracellular domain (ECD). Furthermore, RLKs are known to function as dimers or higher-order oligomers to initiate activation of the intracellular signalling cascade^16^. Homo- and heterodimerization between the ECDs of the receptor or co-receptor partners is responsible for establishing and maintaining specific cellular responses to different stimuli ^20–23^. This allows plant cells to read signals from the local microenvironment ^24^, playing critical roles in plant growth, development, immunity, and symbiosis ^25–27^. Activation of various RLKs in response to different stimuli leads to rapid changes in redox status, mainly but not exclusively, in the apoplastic space. Therefore, discovering the molecular mechanisms of ROS sensing in the apoplast is essential to understand how the information encoded by ROS and other signalling processes is integrated.

A major mechanism by which the cell’s redox status is sensed and regulated is through the oxidation of cysteine residues in proteins ^28^. Cysteines contain a thiol group that reacts with ROS and can be oxidized either reversibly or irreversibly, depending on redox conditions. Proteins with multiple cysteine residues can serve as reversible functional switches where cysteine residues, depending on the ROS levels, can exist either in their reduced state (low ROS) or oxidized state (high ROS). This can directly impact protein localization, stability, and protein binding^29^. The type of oxidatively modifying molecule, amplitude of the ROS response and its duration are crucial factors influencing the action of ROS sensors. The reactivity of cysteines is the key feature of proteins that can act as molecular redox switches ^30^. A well-described example of a redox switch is the redox-dependent transition of NONEXPRESSOR OF PATHOGENESIS-RELATED GENES 1 (NPR1) from an oligomeric (oxidized) to a monomeric (reduced) state ^31,32^. Upon biotic stress, the disulphide bonds that keep NPR1 as an oligomer are reduced, and the active monomeric NPR1 is released to the nucleus to interact with transcription factors and induce the expression of pathogenesis-related (PR) genes ^33^. While most redox switches described in the literature are intracellular proteins, HYDROGEN-PEROXIDE-INDUCED Ca^2+^ INCREASE (HPCA1)/ CANNOT RESPOND TO DMBQ 1 (CARD1), a Leucine-Rich Repeat (LRR) RLK that participates in H_2_O_2_ sensing in the apoplast for stomatal closure as well as quinone perception ^34,35^ was described. This receptor induces intracellular Ca^2+^ increases upon oxidation of a cysteine-rich domain in its ECD after extracellular H_2_O_2_ treatment^35^.

This discovery of the first extracellular ROS sensor raises the question of whether there are additional ROS sensors and how these small and highly reactive molecules are perceived in the extracellular space. The RLK family of Cysteine-rich Receptor-like Kinases (CRKs) has been proposed as promising candidates for ROS sensors in the apoplast due to the characteristics of their ECD ^36^. The CRKs contain multiple cysteine residues in their ECD organized in both conserved (C-X8-C-X2-C), and non-conserved motifs ^37^. Many CRK genes are strongly induced by ROS-generating conditions (e.g. ozone, high light, pathogen attack) and by signals that depend on NADPH oxidases (RBOHs) ^38–40^. Conversely, misexpression of specific CRKs (such as CRK2, CRK5, CRK28, CRK36) alters basal and elicitor-induced ROS bursts, hypersensitive cell death, and systemic acquired resistance^40–45^. CRKs also interact genetically and physically with core immune and cell death regulators (FLS2/BAK1 and RBOHD), influence MAPK activation, and modulate defence hormone pathways (SA/JA/ET), all of which are tightly intertwined with ROS homeostasis ^39,40^. The complex formation with RLKs and other plasma membrane components suggest that homo- and heterodimerization plays a fundamental role in the activation of CRKs and their further signalling ^40,43^. Together, these data support a model in which CRKs sense ROS- or stress-induced changes in apoplastic redox and relay this information through their cytosolic downstream substrates.

Although in the past decade, many studies have linked CRKs to stress-related and developmental processes involving elevated ROS production, sparking extensive speculation about their role in apoplastic ROS sensing, we still lack experimental evidence for the mechanisms of action or the effects of ROS on CRK signalling^15,46,47^. As previously mentioned, for RLKs to become fully active, they need to dimerize or oligomerize in receptor complexes, however, knowledge of oligomerization and interaction partners of the CRK family members remains limited. Additionally, while genetic links have been shown between CRKs and ROS-related processes, there is also limited molecular association between CRK proteins, their interaction partners and the physiological effects of CRK activation.

The current study describes a comprehensive ROS-dependent interaction network involving extracellular domains of the Arabidopsis CRK receptor family. We present this as a resource for the potential landscape of ROS-dependent interactions, which can be refined by filtering expression data to inform hypothesis generation. This study also elucidates a mechanistic association between extracellular cysteine oxidation, receptor dimerization, and the regulation of developmental as well as immune-related signalling pathways. Through the integration of high-throughput interactomics, redox proteomics, structural prediction, and *in planta* functional assays, we demonstrate that specific members of the CRK family, notably CRK28 and its interacting partner CRK17, are good candidates for redox-modulated hubs. These hubs influence processes such as senescence, stress responses, and phenotypes resembling autoimmunity.

## Results

### ROS modulates interactions between CRK family members at the level of extracellular domains

To understand the dynamics of the interplay between CRK family members in the context of ROS perception and signalling, we modified the Cell Surface Interactome (CSI) approach by ^16^ to test the dimerization of CRK ECDs in the presence and absence of ROS, as a modulating molecule. We termed this modified assay ROS-dependent Interactome Assay (RIA^CRK^). We were able to clone the ECDs of 40 out of 44 CRKs from Arabidopsis into bait and prey expression vectors for recombinant protein production in *Drosophila Schneider* S2 cells as described in the Materials & Methods (Supp. Fig. 1, Supp. Table 1)^16,48^. The oxidative environment during protein production negatively affects CRK ECD protein interactions. Therefore, we optimized the CSI protocol to detect CRK interactions by adding a reducing agent, dithiothreitol (DTT), to reset the oxidative status of the ECDs prior to their interaction. The addition of 0.5 mM DTT was optimal for allowing CRK ECDs to interact significantly in a pilot screen (Supp. Fig. 2A, B). Similarly, we assessed the optimal concentration of H_2_O_2_ to test the modulation of interactions. Based on this preliminary screening, we selected 100 µM H_2_O_2_ as the ROS condition for testing in the high-throughput screens (Supp. Fig. 2C). To evaluate whether protein stability is affected by the addition of DTT or H_2_O_2_, we determined the thermal stability of the ECDs. We performed a thermal unfolding assay on CRK17 ECD purified at a larger scale using the baculovirus system in TniH5 insect cells (Supp.Fig. 3A). The melting temperature (T_m_) of CRK17 ECD was not affected by adding 0.5 mM DTT (Sup. Fig. 3 B, C) or by adding 100 µM of H_2_O_2_ (Sup. Fig. 3 D, E), indicating the stability of the ECDs is not compromised by the adjusted experimental conditions used in the RIA^CRK^.

**Figure 1.**
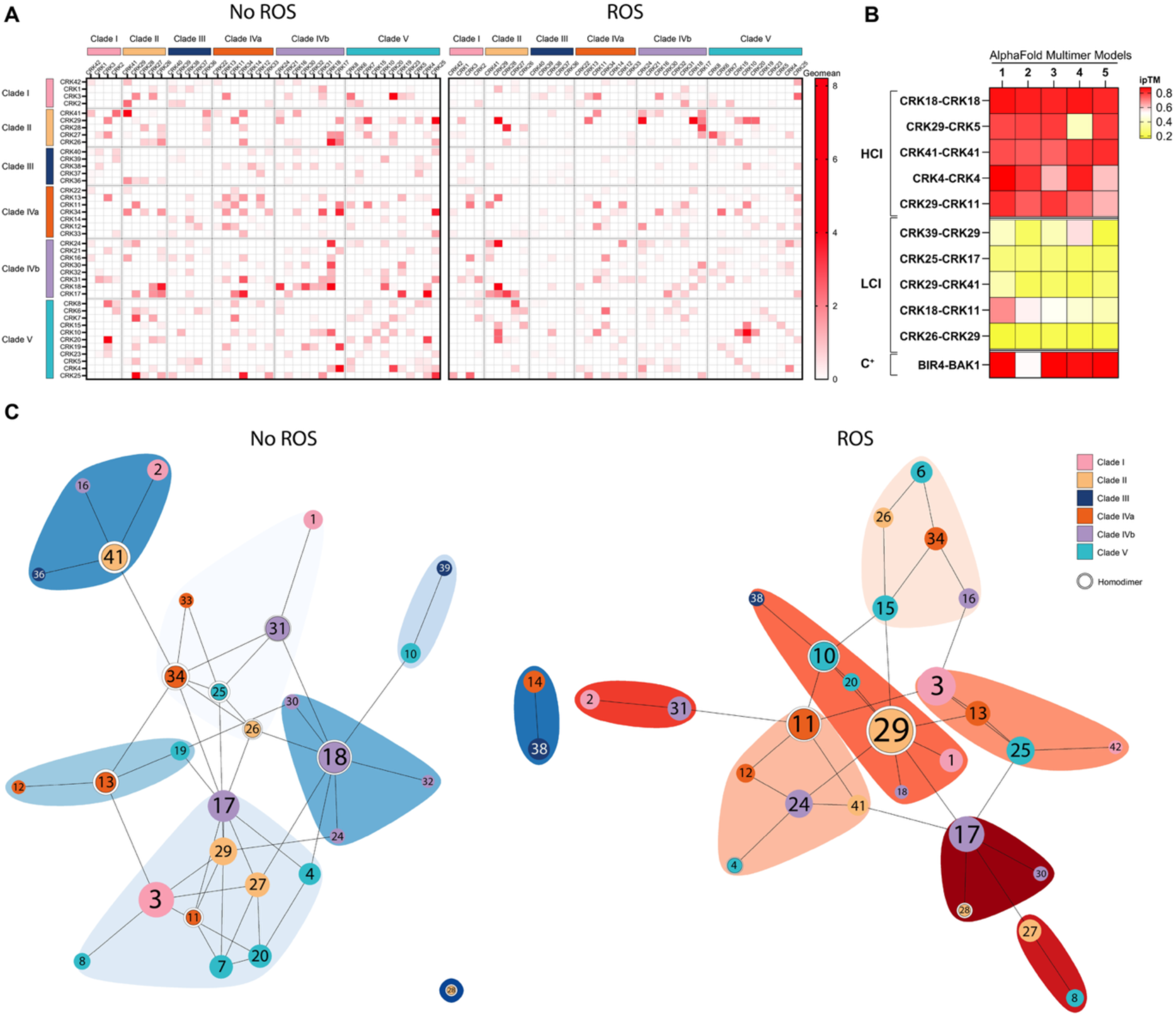
CRK-CRK ECD interactions are modulated by ROS. A) The assembled interaction heatmaps of CRK ECDs for ROS and No ROS conditions, organised by phylogenetic clades as shown in ^36^. The colour scale bar shows interaction scores calculated from the geometric means of modified *Z-*scores as described in Materials & Methods. CRKs are ordered by phylogenetic clade. B) High- and Low-Confidence Interactions from No ROS conditions in A) were modelled using AlphaFold Multimer v2.2.2. The ipTM scores for five predictions per pair are plotted as a heatmap. The scale bar shows ipTM scores from AlphaFold Multimer. C) RIA network of CRK-CRK ECDs interactions for No ROS/ROS conditions. Interactions with GeoMean >1 are plotted. WalkTrap communities are shown in blue or red colours for No ROS/ROS, respectively. Node colours are coded according to CRK phylogenetic clades^36^. A white halo around a node represents a homodimerization interaction. The diameter of the nodes (coloured circles) is proportional to their PageRank score. Numbers in each node correspond to the CRK number (e.g. Node 1 = CRK1). Edges (black lines) show interactions between nodes. RIA networks were generated in No ROS conditions (Schneider’s media with recombinant ECDs diluted 4-fold in TBS) and ROS conditions (same as in No ROS conditions supplemented with an additional 100 µM H_2_O_2_).

**Figure 2.**
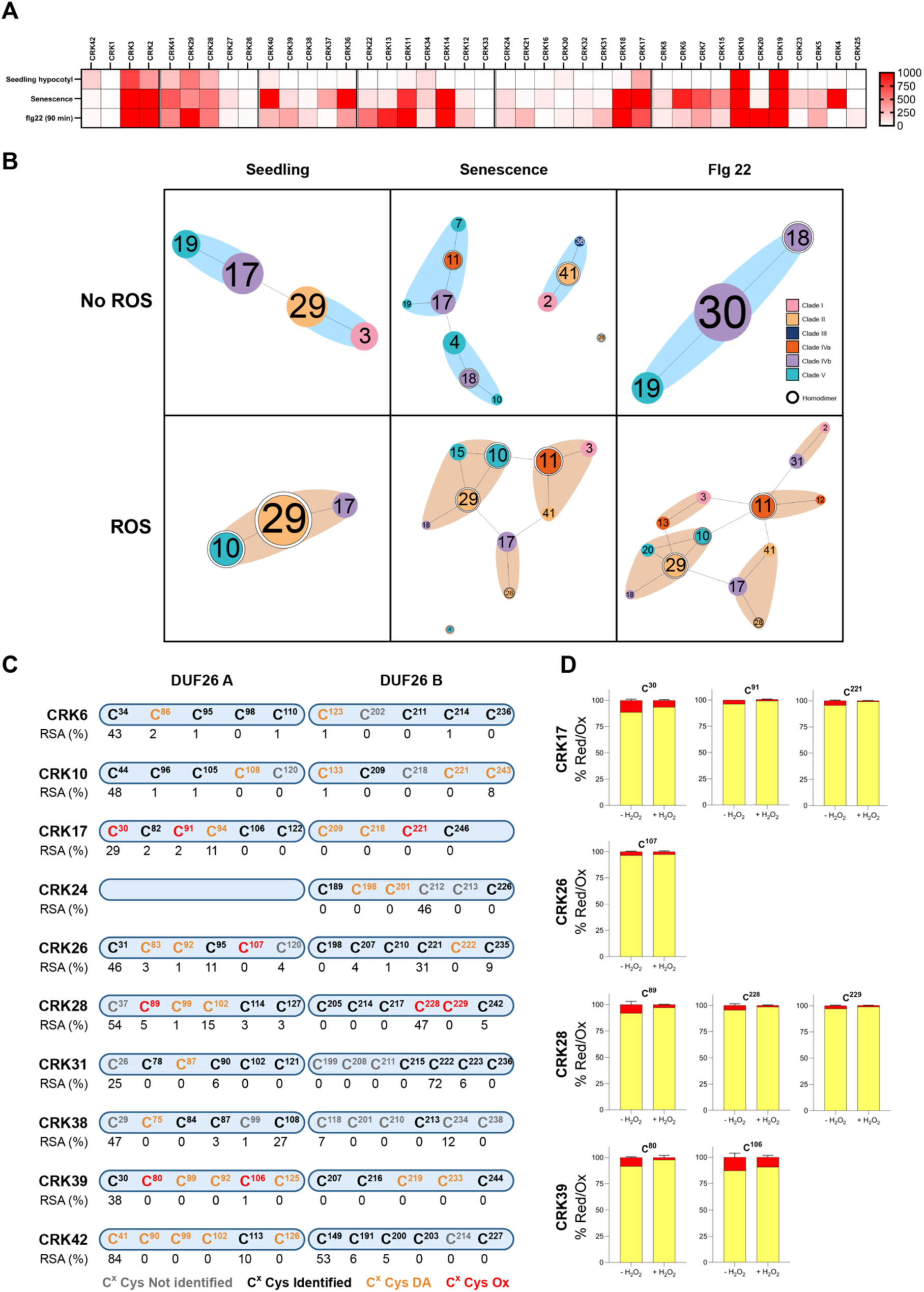
CRKs form ROS-dependent spatiotemporal interaction networks and undergo oxidative modifications. A) Developmental and flg22-triggered atlas of CRK expression. The scale bar represents the absolute counts after DESeq2 mean of ratio normalization. Expression data were mined from TravaDB (Klepikova et al., 2016b) and (Bjornson et al., 2021). B) Spatiotemporal filtered RIA^CRK^ subnetworks from interactions in Fig. 1A. Top panels represent No ROS conditions, bottom panels, ROS conditions. From left to right, filtered by expression in 2-week-old seedlings, senescence, and after 1 µM, 90 min flg22-triggered expression. The No ROS condition in flg22 treatment is after mock treatment, 90 min as per (Bjornson *et al*., 2021). Node colours correspond to phylogenetic clade classification. The white halo around a node indicates homodimerization. Numbers correspond to CRK number (e.g. Node 3 = CRK3). C) Schematic representation of the CRK ECDs identified to be differentially alkylated oxidatively modified cysteines using DAM-MS. CRK ECDs were expressed and secreted from S2 insect cells and treated with PBS (-H_2_O_2_) and 100µM H_2_O_2_ (+H_2_O_2_). Cysteines marked in grey were not identified, in black identified cysteines but not differentially alkylated, in orange identified differentially alkylated cysteines, and in red identified differentially alkylated cysteines that are more oxidized after ROS treatment (reduced percentage of AA, SH). D) Bar plots for the indicated CRKs in which at least one cysteine was found to be more oxidised after H_2_O_2_ treatment.

**Figure 3.**
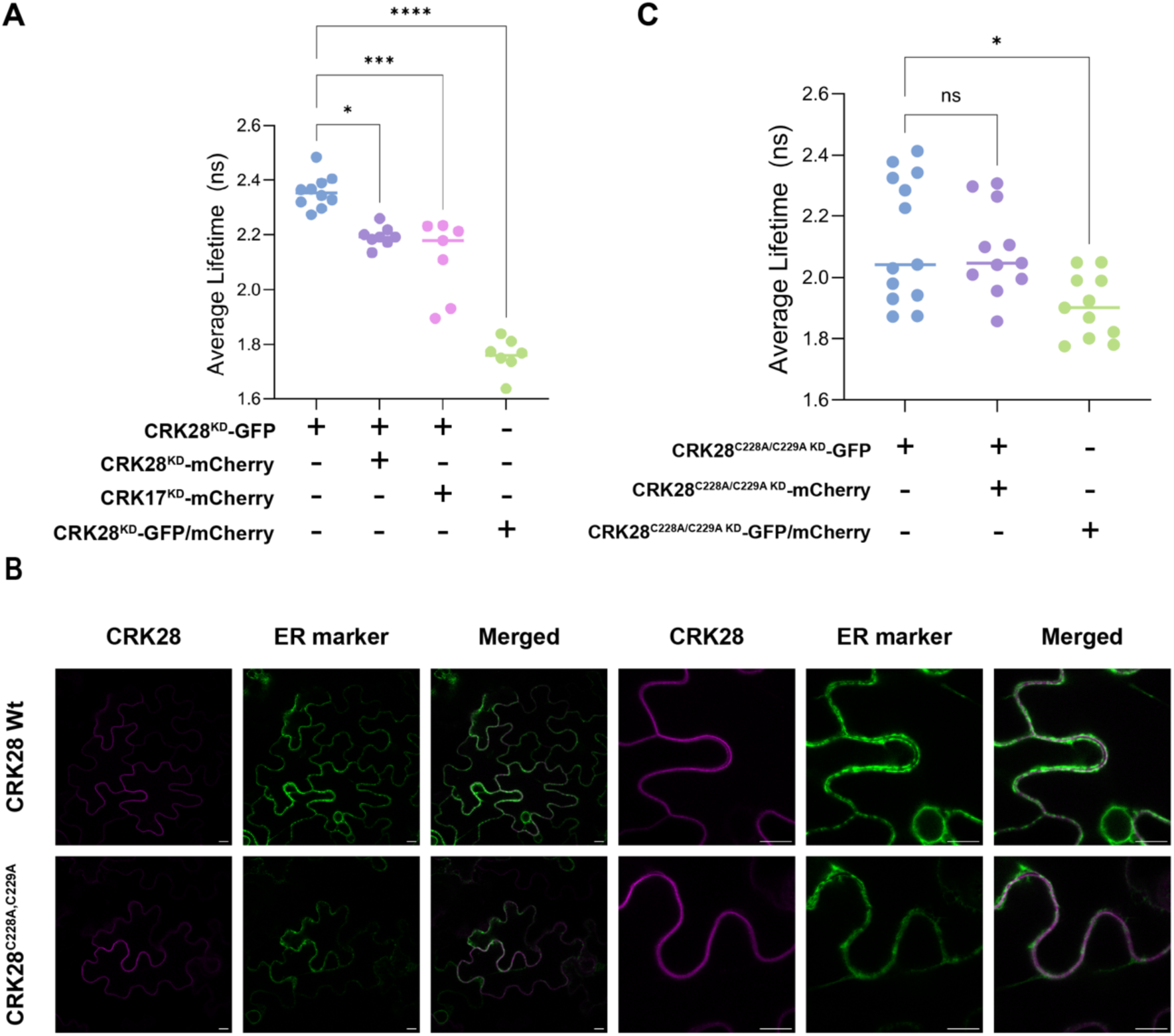
CRK28 homo- and heterodimerizes at the plasma membrane in a cysteine-dependent manner. A) CRK28 homodimerization *in vivo*. CRK28 homodimerization and heterodimerization with CRK17 was measured with FRET-FLIM. B) Transient expression of 35S:CRK28^KD^-3xFLAG-mScI, 35S: CRK28^C228A, C229A-KD^ -3xFLAG-mScI, and 35S:AtVMA21a-mNG (ER-marker) in *N. benthamiana* leaf epidermal cells. Images show an optical section through the cell centre. C) CRK28^C228A, C229A^ mutant homodimerization *in vivo*. CRK28 homodimerization was measured with FRET-FLIM. The average lifetimes in A) and C) were quantified for 3 independent biological repeats. Statistically significant groups were determined by one-way ANOVA and Dunnett’s multiple comparisons test (ns= nonsignificant, **** = p<0.0001). CRK28, CRK17, and the cysteine-to-alanine mutant included the KD mutation.

In the RIA^CRK^ screen, we tested dimeric interactions between bait and prey CRK ECDs under two conditions: No ROS, where no external ROS were added, and ROS, with 100 µM H_2_O_2_ added to the interaction medium. As described before, 40 out of 44 Arabidopsis CRKs ^37,49^ were tested, resulting in screening of 82.82% of all potential interactions. The GeoMean values calculated from reciprocal interactions (Materials & Methods) were plotted as heatmaps in Fig. 1A, in which the CRKs are organized by Clades according to ^36,50^ (Supp. Table 2). In total, 1600 interactions were assessed across two screens, comprising 1560 heterodimeric and 40 homodimeric interactions, for a total of 820 reciprocal pairs. We considered high-confidence interactions (HCI) those with a GeoMean above 1 (Supp. Table 3). Of all tested interactions, only 56/820 and 41/820 were HCI in No ROS and ROS conditions, respectively. Of the HCI found across conditions, 16.07% are homodimers, whereas they represent only 4.8% of all tested interaction pairs, highlighting the relevance of homodimerization to the interactivity of CRK ECDs (Supp. Table 4). Recent advances in structure prediction using AlphaFold ^51^ enabled modelling of CRK ECDs based on crystallized structures of closely related Plasmodesmata Localized Proteins (PDLP5 and PDLP8; PDB: 6GRE, 6GRF)^50^. We used AlphaFold Multimer v2.2.2 ^52]^ to predict dimer structures for five high- and low-confidence interactions detected in the CRK ECDs No ROS interactions screen. Remarkably, all observed HCI were also predicted with high ipTM scores (above 0.75), whereas low-confidence interactions (LCI) were modelled with low ipTM scores for most of the models AlphaFold outputs (Fig. 1B, Supp. Fig. 4, Supp. Table 5).

**Figure 4.**
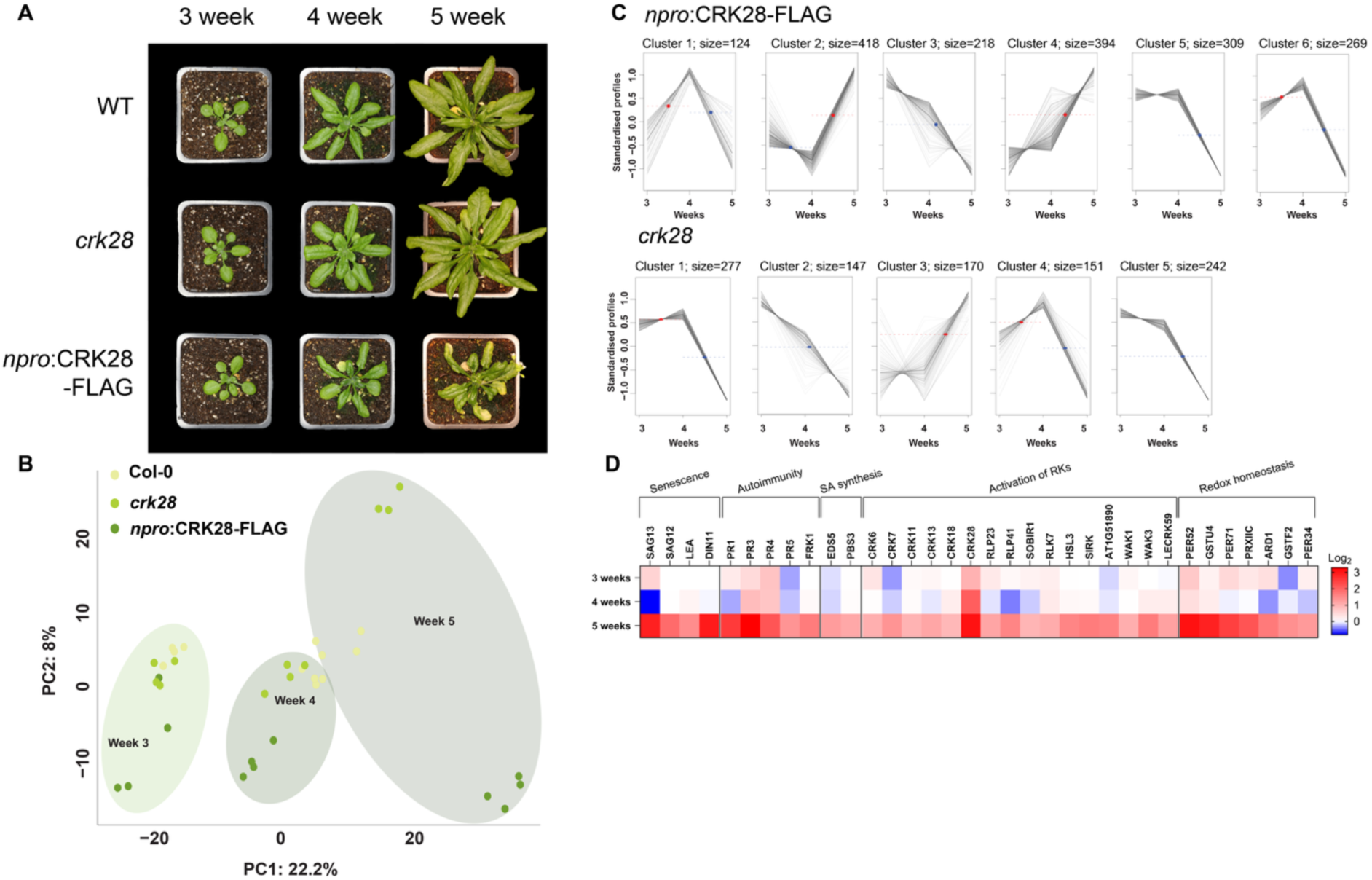
CRK28 functions as a redox-dependent hub receptor. A) 3-, 4-, and 5-week-old Arabidopsis rosettes of representative Col-0, *crk28* T-DNA mutant and npro:CRK28-FLAG lines. B) Principal-Component Analysis (PCA) of the total proteome measurements for all developmental stages (3-, 4-, and 5-week-old plants) and all genotypes indicated in A. C) Time series ordering of all FDR ≥ 0.01 log_2_*Z* scored normalized for *crk28* and npro:CRK28-FLAG in comparison to control Col-0. Clusters are ordered based on the earliest developmental stage. Protein abundance is based on the median time at which all individual profiles in a cluster cross half-maximal abundance within each event window (identified by the red or blue dashed line in the graph and red and blue arrows on the x-axis, respectively). D) A heat map demonstrating changes in protein abundance in the npro:CRK28-FLAG line compared with Col-0 in 3-, 4-, and 5-week-old plants, corresponding to the observed morphological phenotype. Scale bar represents Log2 (Fold Change) of protein abundance.

We constructed a family-wide interaction network of CRK ECDs in the presence and absence of ROS. Only HCI between ECDs were used to construct the network, and communities (groups of highly connected nodes) ^53^ were detected using the walktrap algorithm ^54^ (Fig. 1C, Supp. Table 3) ^53^. This analysis identified eight CRK ECD communities for the No ROS condition and seven for the ROS condition, yielding distinct network architectures for each condition. In the No ROS condition, two very small communities, CRK39-CRK10 and CRK38-CRK14, are disconnected from the main network, whereas the remaining six communities remain cross-connected. In the ROS condition, however, these small, detached networks become integrated into the main body, with CRK10 becoming a more connected, homodimerizing node within one of the central communities. Notably, in the ROS condition, community interconnections are stronger, indicating more interactions between CRK when ROS is present. The nodes in the networks were colored based on the phylogenetic group of each CRK, highlighting emerging interaction patterns among CRK clades. In No ROS, one community with CRK18 as a hub is formed exclusively by members of Clade IVb. However, all the other communities in both conditions are populated by interactions between CRKs across different clades. Based on the HCI score, the top 20 interacting pairs for No ROS are primarily from Clades IVb and V, whereas after ROS treatment, they shift to Clades II and V (Supp. Table 6). In particular, the HCI of interactions between two closely related homologues, CRK28 and CRK29, accounts for most of the top 10 interactions under ROS conditions.

Using page rank and betweenness as measures of node centrality, we identified network hubs (Supp. Table 7). These hubs can be understood as highly-interacting ECDs crucial to the regulatory network controlled by CRK ECDs ^55^. Under no ROS conditions, CRK17/18/29/34/41 serve as main hubs, whereas in ROS conditions, CRK17/29 remain hubs, CRK10/11/15 increase in centrality, and CRK18 is no longer a hub. The condition-dependent network architecture confirms that CRK ECD interactomics is ROS-sensitive, pointing to a likely ROS-sensing function of the CRK machinery.

### Spatiotemporal CRK interaction networks

The RIA^CRK^ screen represents the potential landscape of CRK ECD interactions modulated by the presence or absence of H_2_O_2_. However, to assess whether the interactions identified in the *in vitro* RIA^CRK^ screen are possible *in vivo*, we filtered the interactome using the expression data of *CRK* genes. We retrieved the expression of CRKs in whole seedlings and senescent leaf tissue from TravaDB ^56^ and in 2-week-old plants treated for 90 min with 1µM flg22 from the work of ^57^ as Absolute counts (Fig. 2A, Supp. Table 8). In these developmental stages, the expression patterns of CRK family members differ substantially for each CRK member. We then filtered the RIA^CRK^ No ROS and ROS conditions using the expression data to show spatiotemporal-filtered networks for seedlings, senescent leaves, and seedlings treated with flg22 (Fig. 2 B). Treatment with flg22 leads to rapid production of ROS in the apoplast; therefore, we filtered the No ROS by using the expression of CRKs in mock-treated plants instead of flg22-treated ones, for a better representation of physiological conditions. The interaction networks are modulated in two dimensions, depending on whether ROS is present in the interactome and whether CRKs are expressed in the specific developmental stage or upon treatment (Fig. 2B). Nodes connected by edges were plotted only if both genes were considered expressed in the tissue (>195 absolute units). The size of seedling networks is dramatically decreased due to the low expression of most CRKs in seedlings. Under ROS conditions, node identities remain unchanged, but interaction dynamics shift, with CRK29 becoming the central hub in the ROS network. Interestingly, we observed that the senescence network is substantially larger than that of seedlings (Fig. 2B). As senescence is characterized by an enhanced production of ROS, the senescence-filtered No ROS subnetwork might not represent a real physiological state. However, in the presence of ROS, the communities are highly connected, with most nodes connecting to two or three other CRKs, except for CRK4 homodimerization. Flg22 also elevates the expression of many CRKs, as seen in the subnetwork composition. In the mock-treated and No ROS conditions, we observe minimal CRK expression, similar to that in seedlings, with only CRK18/19/30 expressed and interacting. On the other hand, under ROS conditions, the composition of the communities is highly similar to the senescence-filtered one, with high interconnection and presence of CRK11 and CRK29 as hubs. Overall, these spatiotemporal subnetworks offer clearer insights into potentially physiologically relevant interactions within ROS-modulated signalling cascades.

### Mapping of Redox-Sensitive Cysteines in CRKs

To determine whether cysteines in CRK ECDs can undergo oxidative modifications, we use the Differential Alkylation Method (DAM) followed by mass spectrometry (MS) measurement (DAM-MS). With this technique, the oxidation state of cysteine thiol groups is detected via sequential alkylation with alkylation agents. First, reduced thiols (-SH) are alkylated by acrylamide (AA), blocking them from further modification. Then, the remaining thiols reversibly oxidized as disulphide bonds (S-S) or as sulfenic acid (SOH) are reduced and subsequently alkylated with iodoacetamide (IAM). The ratio between AA- and IAM-tagged peptides can be calculated as an “oxidation percentage” for each cysteine (Supp. Fig. 5A). To assess the dynamics of cysteine oxidation upon ROS treatment, we performed DAM followed by MS on CRKs ECDs in No ROS and ROS conditions, as in RIA^CRK^. These ECDs were cloned into an S2 cell expression vector with the V5 tag lacking the Fc Fragment. In this conformation, only 32 CRK ECDs had sufficient expression to perform DAM (Supp. Fig. 5B).

**Figure 5.**
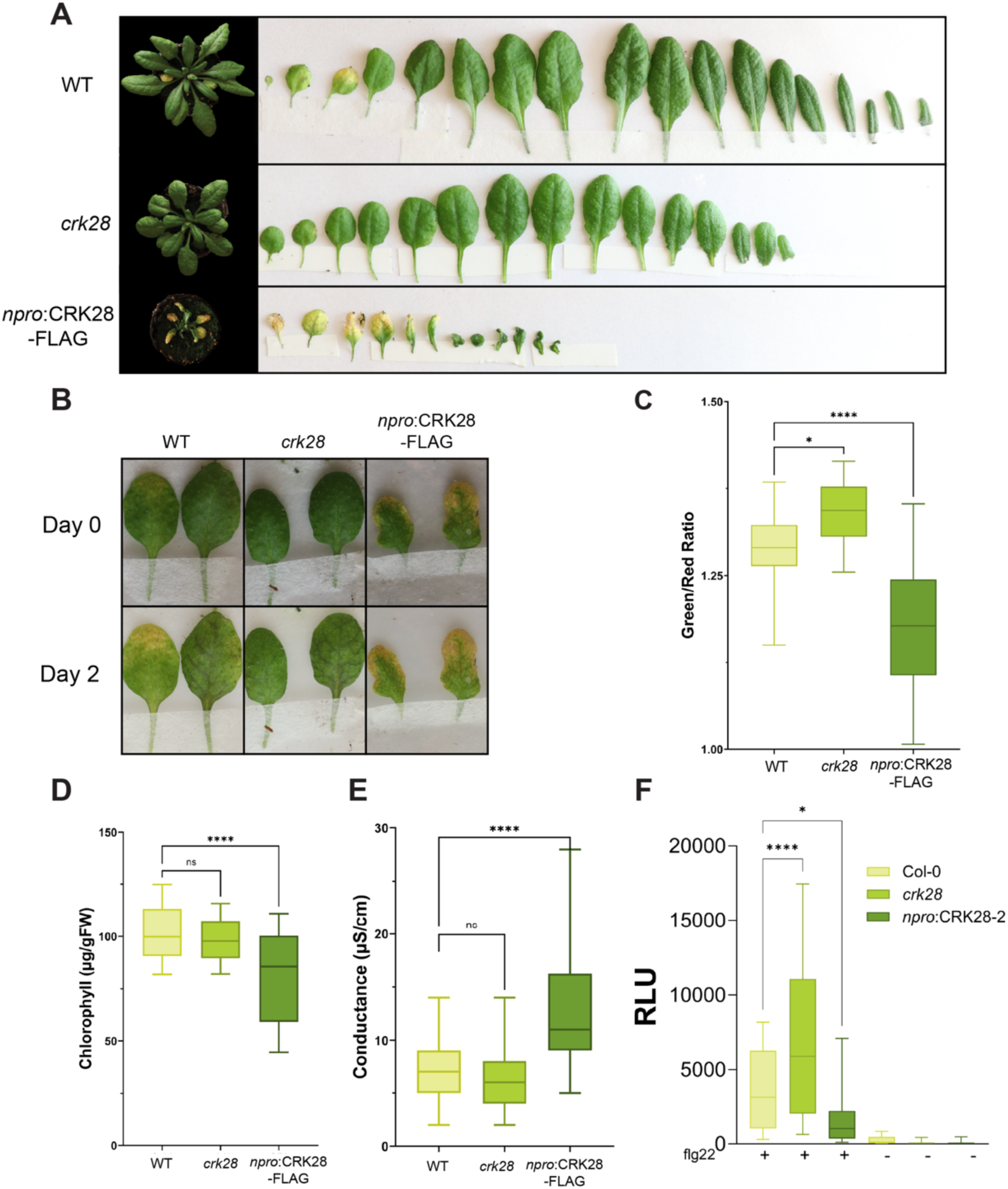
CRK28 as a regulator of senescence and an autoimmunity-like state. A) 5-week-old Arabidopsis rosettes and leaf panels of representative Col-0, *crk28* T-DNA mutant and *npro*:CRK28-FLAG lines. B) Leaf detachment assay on the 5^th^ and 6^th^ leaves of 5-week-old plants. Pictures taken at Day 0 and Day 2 to observe senescence progression. Representative images of samples of 5^th^ leaf (left) and 6^th^ leaf (right) in each square. D) Quantification of green-to-red ratio pixels from pictures in B) with ImageJ plugin GreenLeafVI. E) Chlorophyll content measurement of 4 mm leaf disks of the 7^th^ and 8^th^ rosette leaves of 5-week-old plants. F) Ion leakage measurements of leaf disks in D. G) Sum of luminescence of ROS burst curves on 4 mm leaf disks of the 9^th^ and 10^th^ rosette leaves of 5-week-old plants triggered by 1 µM flg22 peptide. Asterisks indicate statistical significance in a one-way ANOVA for each sample compared to Col-0. ns= not significant, * p-value < 0.5, **** p-value <0.0001. Error bars represent standard deviations in D-G, N=24 in all experiments.

*In silico* prediction showed that out of the 360 cysteines present in the 32 CRK ECDs produced, only 78.3% (282) cysteines are measurable, as the size and properties of certain peptides generated after trypsin digestion make the remaining 21.7% impossible to detect by MS (Supp. Fig. 5C). Therefore, 133 out of 282 measurable cysteines were detected in the 32 CRK ECDs. The distribution of identified cysteines varied substantially across CRK ECDs (Supp. Fig. 5D). This depends on several factors, including the frequency of trypsin target sites in the protein sequence, the efficiency of peptide digestion and labelling, and partial reduction and denaturation during DAM. The number of differentially alkylated (DA) cysteines in relation to the total number of cysteines measured per CRK ECD is shown in Supp. Fig. 5E and Supp. Table 9. The number of DA cysteines rarely exceeds 50% of the total identified cysteines, and most frequently, only 2-3 DA cysteines are identified (Fig. 2C, D Supp. Fig.5 E). Overall, 10 out of the 32 CRK ECDs tested underwent differential labelling with the alkylation agents, as shown in Fig. 2C. Out of these, 4 of them (CRK17/26/28/39) showed more than 2% higher oxidation after treatment with H_2_O_2_, in at least two independent repeats per condition (Fig. 2C and D, Supp. Fig.5E).

Crystal structures of PDLPs and AlphaFold CRK models predict that most cysteine residues in CRK ECDs are likely involved in disulphide bonds ^58^. Based on the stability of cysteines in PDLP ECDs, it was hypothesized that cysteines forming disulphide bonds in CRKs are structural and therefore important for the folding of the protein, but not functional as redox switches^50^. However, several CRKs from Clades II, III and IV have extra non-conserved cysteines compared to PDLPs. Additionally, CRKs in Clades II and IV have sequentially adjacent (also termed vicinal) cysteines, like CRK28 with C228 and C229 (Supp.Fig.5F). To perform ROS sensing through oxidative modification, these residues should be solvent-accessible; therefore, we calculated the relative solvent-accessible surface area (RSA) in PyMol as a measure of solvent exposure for each DA cysteine of the 10 CRK ECDs in Fig. 2C. RSA is defined as the solvent-accessible area relative to the maximum solvent-accessible area for each residue ^59^. Some of the significantly oxidized cysteines, such as CRK17 C30 and CRK28 C228 are predicted to have high RSA values (Fig.2C, Supp. Table 9). Given the presence of vicinal cysteines, high RSA, and elevated oxidation after ROS treatment, we hypothesize that C228 and C229 of CRK28 can function as redox switches.

### CRK28 homodimerizes and heterodimerizes with CRK17 *in vivo*

CRK28 emerges as a promising candidate for an ROS sensor via direct oxidative modification of cysteines in its ECD. Although CRK28 homodimerizes in both No ROS and ROS conditions in the RIA^CRK^, the interaction is stronger upon ROS treatment *in vitro* (Supp. Figure 6). It also interacts with CRK17 in ROS conditions, and both of them have cysteines found to be significantly oxidized *in vitro* (Fig. 2C, D, Supp. Figure 6). Additionally, CRK28 expression is induced upon biotic stress and senescence, two physiological stages characterized by elevated ROS production.

**Figure 6.**
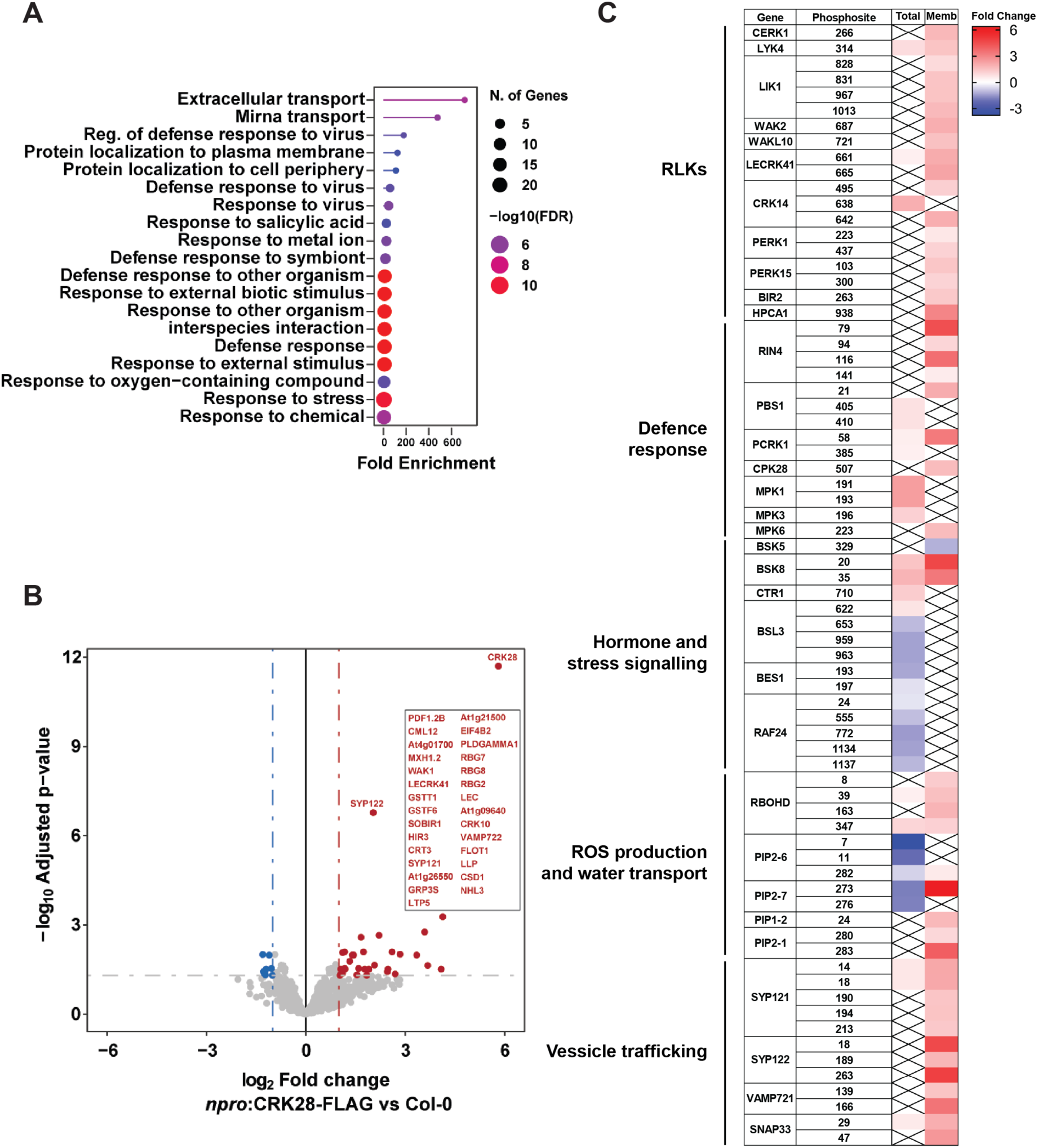
CRK28 interacts with stress-related partners and, upon overexpression, rewires the phosphoproteome in 5-week-old plants. A) GO enrichment for statistically significant interacting partners of CRK28 identified using IP-MS. B) Volcano plot of IP-MS of 5-week-old Arabidopsis rosettes of the npro:CRK28-FLAG line compared to Col-0. In volcano plots, horizontal dashed lines correspond to 0.05 FDR, and vertical lines correspond to -1 and 1 Log2 (Fold change) as statistical cutoffs. The important interacting partners are marked. C) Heatmap of significant hyper- and hypophosphorylated proteins in the total extract and membrane-enriched phosphoproteomes of 5-week-old Arabidopsis rosettes of npro:CRK28-FLAG line compared to Col-0. Complete list in Supp. Tables 12 and 13.

To confirm that CRK28 dimerises *in vivo* on the level of the full-length receptor, we used Förster Resonance Energy Transfer - Fluorescence-Lifetime Imaging Microscopy (FRET-FLIM) to determine CRK28 homodimerization and heterodimerization with CRK17. FRET-FLIM measures the decrease in the lifetime of a donor fluorophore (GFP) due to energy transfer to an acceptor (mCherry) when they are in close proximity to each other. It was previously reported that kinase-dead point mutations of CRK28 bypass the induced cell death processes resulting from CRK28 overexpression in Arabidopsis and *Nicotiana benthamiana*^40^. Substituting the corresponding conserved lysine in the ATP-binding site of the kinase domain for asparagine in CRK28 and CRK17 (K377N and K387N, respectively) increased expression levels. CRK28 and CRK17 with kinase-dead point mutations were expressed transiently in *Nicotiana benthamiana* under an estradiol-inducible promotor (CRK28^KD^-GFP, CRK28^KD^ -mCherry, and CRK17^KD^ -mCherry). The fluorescence timeline of CRK28^KD^-GFP at the plasma membrane decreased in the presence of both CRK28^KD^ -mCherry and CRK17^KD^ -mCherry acceptors, from 2.40 ± 0.04 ns to 2.20 ± 0.18 ns (Fig. 3A). This indicates that CRK28 is able to homodimerize but also to heterodimerize with CRK17 at the level of the full-length receptor.

### Cysteines 228 and 229 play a functional role in CRK28

To investigate the role of cysteine residues in CRK28, we assessed the effects of cysteine mutants on the localization and functionality of CRK28 *in vivo*. We generated double cysteine mutant substitutions to alanine for each of the six predicted disuphide bonds of CRK28 (CRK28^C37A,C114A^, CRK28^C89A,C99A^, CRK28^C102A,C127A^, CRK28^C205A,C214A^, CRK28^C217A,C242^ and CRK28^C228A,C229A^). All cysteine to alanine substitutions were introduced, as for the FRET-FLIM studies, to the kinase dead (KD) version of CR28. CRK28^WT^ and cysteine mutants were transiently expressed in *Nicotiana benthamiana* under a constitutive promoter and transcriptionally fused to a fluorescent tag (35S:CRK28^KD^-3xFLAG-mScI). The CRK28^WT^ and CRK28^C228A, C229A^ variants localize to the plasma membrane, whereas the other CRK28 cysteine mutants show a patchy expression pattern similar to the endoplasmic reticulum (ER)-marker, suggesting they are retained in the ER (Fig.3B, Supp. Fig. 7 and 8). This suggests that most cysteines in DUF26A and B form structurally important disulphide bonds required for the proper folding and localization of the proteins, while C228 and C229 are not involved in protein localization.

To test whether C228 and C229 are important for the function of CRK28, we tested the homodimerization capability of the double cysteine mutant using the previously described FRET-FLIM system (Fig. 3C). We observed that the CRK28^C228A,C229A^-GFP construct has a lower fluorescent lifetime on its own compared to the CRK28^KD^-GFP, however, it does not decrease further when co-expressed with the acceptor. This indicates that the cysteine mutant is unable to homodimerize. We further confirm the redox reactivity of these cysteines *in vivo* by measuring oxidative modifications in CRK28 ECDs after transient expression in *Nicotiana benthamiana*. We identified C228 as significantly oxidatively modified after H_2_O_2_ treatment (Supp. Fig. 9). In light of these results, it appears that C228 and C229 are important for the biological function of CRK28 (Supp.Fig.10).

### CRK28 is important for plant development

To investigate the role of CRK28 in plant development, we evaluated whether the *crk28* knockout line (crk28 T-DNA; SALK_085178) and a complementation line of CRK28 driven by the native promoter in the *crk28* T-DNA background, npro: CRK28-FLAG, exhibit any significant morphological abnormalities when compared to Col-0 plants^40^. These genotypes were grown under optimal conditions (long-day: 16 hours of light and 8 hours of darkness at 21 °C). At the two-week-old seedling stage, we observed that the *crk28* line exhibited an increase in primary root growth of approximately 10-15% compared to wild-type Col-0 control plants (Supp. Fig. 11), as reported earlier by Pelagio-Flores et al. (2020). However, plants expressing CRK28 under its native promoter, npro: CRK28-FLAG, were indistinguishable from Col-0 in terms of primary root length. While tracking the development of these lines over time, it became apparent that the rosettes of line npro: CRK28-FLAG began to exhibit a slightly dwarf phenotype at the stage of 4-week-old rosettes. In this line, we also observed earlier and more pronounced senescence symptoms than in *crk28* and Col-0 plants (Fig. 4A). These morphological changes were substantially intensified in 5-week-old npro: CRK28-FLAG plants. At this stage of development, we observed a 15-fold higher mRNA expression in npro: CRK28-FLAG compared to wild-type (Supp. Fig. 12A). At the stage of 5-week-old plants, the leaf number and morphology of 5-week-old *crk28* and npro: CRK28-FLAG lines are different from those of Col-0 plants (Fig. 5A). In Col-0, the first fully expanded juvenile leaves are rounder and have smoother edges, while mature leaves are more elongated and have serrations in the margin ^60,61^. The transition from juvenile to mature leaves occurred consistently in Col-0 between the 5th and 6th leaf. The *crk28* mutant plants showed a delayed transition to mature leaves, occurring between the 6th and 7th leaf, and maintained a broader shape compared with the elongated Col-0 mature leaves. On the other hand, the npro: CRK28-FLAG plants transitioned from juvenile to mature leaves earlier, and the mature leaves were extremely stunted and curled. Additionally, we also observed significant yellowing of the juvenile leaves in the npro: CRK28-FLAG line. This indicates that npro: CRK28-FLAG plants show early signs of senescence at 5 weeks.

To understand the molecular basis of these morphological changes, the total proteome was measured in the *crk28* and npro: CRK28-FLAG in 3-week-old young plants, 4-week-old adult plants before transitioning to the generative stage, and 5-week-old plants at the onset of leaf senescence. We could not detect a major difference between *crk28* and npro: CRK28-FLAG in comparison to Col-0 plants at the stage of 3-week-old plants and only very marginal at the stage of 4-week-old plants (Fig. 4B). However, for 5-week-old plants, we measured a substantial difference in protein abundance for *crk28* and npro: CRK28-FLAG in comparison to Col-0 plants. We also measured the progressive accumulation of the CRK28 protein in 3-, 4-, and 5-week-old rosette leaves of the npro:CRK28-FLAG line. This confirms the overaccumulation of CRK28 during later developmental stages compared to Col-0 plants (Supp.Table 10).

Clustering of differentially expressed proteins in *crk28* and npro: CRK28-FLAG was performed based on their expression patterns throughout development (Fig. 4C). This resulted in five and six unique clusters for c*rk28* and npro: CRK28-FLAG, respectively. Gene ontology enrichment was used per cluster to identify biological processes affected in the *crk28* and complementation line (Supp. Fig.13). Interestingly, in the *crk28* line, proteins mainly linked to protein synthesis (clusters 1, 4, and 5) and secondary metabolism (clusters 2 and 3) showed differential expression compared with Col-0 plants. This suggests that many developmental processes are ongoing, either to maintain the vegetative stage or to prepare for the shift from the vegetative to the generative phase. However, in the npro: CRK28-FLAG line, there is a gradual increase in proteins associated with stress responses, including defence, systemic acquired resistance, senescence, and redox homeostasis (clusters 2 and 4). Conversely, proteins crucial for photosynthesis and chloroplast integrity (clusters 1, 3, 5, and 6), as well as those involved in protein synthesis (clusters 5 and 6), decline over time (Supp. Fig.13).

In fact, in the proteomic analysis we identified elevated levels of the senescence protein markers, such as Senescence-Associated Genes 12 and 13 (SAG12 and SAG13), Late Embryogenesis Abundant (LEA) and Dark-Inducible 11 (DIN11) in npro: CRK28-FLAG line at the stage of 5 weeks (Fig. 4D). The *crk28* mutant showed delayed senescence compared to Col-0 quantified by the green-to-red pixel ratio in the pictures taken at two days post-detachment, while the npro: CRK28-FLAG line showed a much more advanced yellowing already at Day 0 and faster progression at Day 2 (Fig. 5B, C). Also, plants overexpressing CRK17 display early onset of senescence (Supp. Fig.12). The chlorophyll content measured for *crk28* was not significantly enhanced compared to Col-0, but it was reduced considerably in the npro: CRK28-FLAG line (Fig. 5D). Similarly, programmed cell death, which is a late-stage hallmark of senescence, was more pronounced in the leaves of the npro: CRK28-FLAG line, but not in *crk28* (Fig.5E). These results suggest that CRK28 is a regulator of processes related to the initiation of leaf senescence.

On the other hand, the phenotype of the 5-week-old npro: CRK28-FLAG line with stunted growth, dwarfism and widespread necrotic lesions on leaves, also strongly resembles the autoimmunity mutant phenotype^62–64^. Alongside plant ageing and the accumulation of CRK28, we observed elevated levels of protein associated with constitutively active defence responses: such as Pathogenesis-Related (PR) members PR-1, 3, 4, and 5, as well as FLG22-induced receptor-like kinase 1 (FRK1)/ Senescence-induced receptor-like serine/threonine-protein kinase (SIRK) (Fig.4D). Moreover, increased expression of the enzymes involved in salicylic acid (SA) synthesis and the constitutive activation of plasma membrane receptor kinases or receptor-like proteins from families such as CRKs, LRR-RKs, WAKs, or RLPs were also observed as characteristic features of plant autoimmunity (Fig.4D). Having observed enhanced autoimmunity markers along redox homeostasis-related proteins in line with elevated CRK28 dosage, we next wanted to assess the ability of these plants to mount a canonical defence response. To do so, we tested the flg22-triggered ROS burst in 5-week-old *crk28* and npro: CRK28-FLAG plants, compared with Col-0 (Fig.5F). ROS production was reduced in npro: CRK28-FLAG plants, whereas in the knockout line it was elevated compared to Col-0. This all suggests that the increased abundance of CRK28 in the npro: CRK28-FLAG plants at the late developmental stage, when its endogenous level is already elevated, creates conditions in which ROS levels might already be high, leading to accelerated senescence and autoimmune responses, in which flg22-triggered ROS burst is attenuated.

### CRK28 associates with stress-related proteins to regulate ROS-signalling

Immunoprecipitation followed by MS was used on 3, 4 and 5-week-old npro: CRK28-FLAG plants to assess CRK28 complexes at the plasma membrane (Fig. 6A and B, Supp. Table 11). No significant interactors were observed at weeks 3 and 4, likely due to low CRK28 expression (Supp. Table 11). However, proteome data in week 5 showed 62 significant interactors (Figure. 6B, Supp. Table 11). GO term enrichment revealed that these proteins can be classified into stress response proteins and extracellular transport (Fig.6A). We identified several PR family members that mainly enhance resistance to biotic pathogens (fungi, bacteria, viruses, nematodes) and respond to abiotic stress ^65–67^. The most confident interactor was plant defensin 1.2b (PDF1.2b), a cationic antimicrobial peptide offers broad pathogen resistance, especially against necrotrophic fungi ^68,69^. We also detected Lipid transfer protein 5 (LTP5, PR-14), which breaks down microbial membranes, and Pathogenesis-related gene 5 (PR-5), which is important for antifungal activity. Moreover, we identified several proteins, like glutathione S-transferase THETA 1 (TGSTT1), copper-zinc superoxide dismutase 1 (SOD1), Glutathione S-transferase F6 (GSTF6), and Peroxidase 67, involved in the removal of excess ROS from the extracellular space and known to be redox-regulated via changes in the oxidation status of cysteine residues^70,71^. All these proteins are secreted into the extracellular space and are mainly plant antioxidant defence proteins involved in redox regulation within plant cells.

CRK28 associates with plasma membrane receptors such as Cell Wall-Associated Kinase 1 (WAK1), which connects the cell wall pectin matrix to the cytoplasm, and is crucial for cell wall integrity, cell expansion, and immune responses. Additionally, we identify the Suppressor of BIR1 (SOBIR1), which acts as a co-receptor for Leucine-rich repeat receptor-like proteins (LRR-RLPs) to mediate immune responses against diverse pathogens, and also interacts with CRK28. Interestingly, SOBIR1 functions as a checkpoint that prevents autoimmunity. Overexpression of SOBIR1 or disruption of its negative regulator, BIR1, leads to constitutive activation of defence responses and cell death^72–74^. This aligns well with the morphological phenotype observed for 5-week-old npro: CRK28-FLAG plants (Fig. 5A).

The association of PR and ROS detoxification proteins, selected RKs, and CRK28 is consistent with the parallel detection in the same pool of interactors the (SYNTAXIN OF PLANTS 121 / PEN1) SYP121, (SYNTAXIN OF PLANTS 122) SYP122, and (VESICLE-ASSOCIATED MEMBRANE PROTEIN 722) VAMP722 proteins^75,76^. SYP121 and SYP122 (Qa-SNAREs) and VAMP722 (R-SNARE) form complexes that mediate vesicle fusion at the plasma membrane, a process implicated in the targeted secretion of defence-related cargo, including PR proteins and ROS-modulating enzymes, to the apoplast. Their co-enrichment therefore supports a model in which SNARE-dependent vesicle trafficking coordinates ROS-associated processes.

The consequence of a complex formation event is most often the activation of signalling cascades via phosphorylation. Therefore, we examined the phosphoproteome of whole-cell fraction and membrane-enriched samples of *crk28* and npro: CRK28 compared with Col-0 at the stage of the 5-week-old plants with morphological phenotypes (Fig.4A). In the total and membrane phophoproteomes there were approximately tenfold more hyper- and hypophosphorylated peptides identified in the npro: CRK28-FLAG than found in *crk28* plants (Supp.Fig 14A and B, Supp. Tables 12-17). The wide variety of differently phosphorylated peptides in the npro: CRK28-FLAG plants indicates a broad reorganization of the cellular phosphoproteome at week 5, a time when the plants experience morphological changes likely caused by increased CRK28 levels. Based on the GO term enrichment in *crk28* plants, we identified mostly changes in the phosphorylation of proteins related to photosynthesis-related processes, as well as protein localisation and transport, and protein synthesis (Supp.Fig. 14C). However, npro: CRK28-FLAG plants modulated phosphorylation mostly occurs for the proteins related to stress response and vesicle trafficking biological processes (Supp.Fig 14C). Additionally, active kinases identified in both lines show more pronounced or opposite phosphorylation status in the line overexpressing CRK28 compared to the *crk28* mutant (Supp.Fig.14C).

The enhanced recruitment of the pattern-recognition and defence-related receptors to the plasma membrane at the protein level is accompanied by the activation of their kinase domains. We identify several differentially phosphorylated RKs, including Chitin elicitor receptor kinase 1 (CERK1), LysM domain receptor-like kinase 4 (LYK4), LysM RLK1-interacting kinase 1 (LIK1), Wall-associated receptor kinases 2 and 10 (WAK2 and WAKL10), L-type lectin-domain containing receptor kinase IV.1 (LECRK41), Proline-rich receptor-like protein kinase PERK1 and 15 (PERK1 and 15) and BAK1-INTERACTING RECEPTOR-LIKE KINASE 2 (BIR2) in npro: CRK28-FLAG (Fig.6C). Among these defence-related receptors, we also observed HPCA1, an extracellular H_2_O_2_ sensor, with enhanced phosphorylation at T603, S916, and S938, residues that had not previously been reported to undergo autophosphorylation after H_2_O_2_ treatment. This may indicate that the observed hyperphosphorylation is attributable to interactions with other kinases, consequent to elevated CRK28 levels.

Along with this extensive activation of plasma membrane receptors, other immune regulators such as RIN4, AvrPphB susceptible protein 1 (PBS1), PTI-COMPROMISED RECEPTOR-LIKE CYTOPLASMIC KINASE 1 (PCRK1), and Calcium-dependent protein kinase 28 (CPK28) were also hyperphosphorylated in npro: CRK28-FLAG but absent in *crk28* (Fig.6C, Supp. Table 12 and 13) ^77–85^. We observe hyperphosphorylation of the MAP kinases MPK1 (Thr191 and Thr193), MPK3 (Thr196), and MPK6 (Tyr223), which function in developmental and stress processes ^86,87^. Components of brassinosteroid and ethylene signalling were also affected, including Brassinosteroid-signalling kinase 5 and 8 (BSK5 and BSK8) and CONSTITUTIVE TRIPLE RESPONSE1 (CTR1)^88^. In parallel, many kinases were hypophosphorylated, among them BSU1-like proteins 2 and 3 (BSL2 and BSL3), BRI1-EMS-SUPPRESSOR 1 (BES1), or RAF-like kinase 24 (RAF24). RAF24 dephosphorylation on multiple residues is very interesting, as RAFs are critical Mitogen-Activated Protein Kinase Kinase Kinases (MAPKKKs) that function as central integrators for environmental stress responses (e.g., drought, salinity) and growth regulation (Fig.6C) ^89^.

Interestingly, we found that NADPH oxidase RBOHD, a key component in signalling for immune responses and environmental stress tolerance, showed multiple hyperphosphorylated residues (8, 39, 163, 347), most of which, except for 8, are associated with the activation of defence responses. ^90,91^. Activation of the RBOHD as a main hub that generates apoplastic ROS leads to differential phosphorylation of Aquaporins (AQPs), specifically Plasma Membrane Intrinsic Proteins (PIPs), which play a crucial role in regulating plant growth, development, and stress responses by facilitating the transport of ROS, particularly hydrogen peroxide, across cellular membranes^75,92^. PIPs enable apoplastic H2O2 to enter the cytoplasm, transducing external signals into cellular responses. We found that PIP2-6 and PIP2-7 are hypophosphorylated, while PIP1-2 and PIP2-1 are hyperphosphorylated (Fig. 6C).

Lastly, SNARE components SYP121, SYP122, VAMP721 (all of these were found to interact with CRK28), and SNAP33 were strongly phosphorylated, indicating extensive reconfiguration of plasma membrane trafficking ^93–95^.

Together, these data demonstrate that CRK28 overaccumulation is associated with broad rewiring of phosphorylation networks affecting immune signalling, ROS production, vesicle dynamics, hormone pathways, and membrane transport, particularly at the plasma membrane, consistent with the autoimmune-like and accelerated senescence phenotypes observed in 5-week-old plants.

## Discussion

This work uncovers a ROS-dependent interaction network among Arabidopsis Cysteine-Rich Receptor-Like Kinases. It highlights a mechanistic connection between extracellular cysteine oxidation, receptor dimerization, and the regulation of ROS signalling involved in development and immune responses. By integrating a high-throughput CRK ECD redox interactome assay, redox proteomics, and structural prediction, we pinpoint several CRK receptors that may act as transducers of ROS signalling. Based on this assay, we use *in planta* functional studies to demonstrate that one of those, CRK28, serves as a redox-regulated hub influencing senescence, stress responses, and autoimmune-like phenotypes.

### ROS remodels the CRK interaction landscape at the ECD level

The RIA^CRK^ assay demonstrates that CRK extracellular domains form dense, selective interaction networks in which only about 5% of the theoretically possible dimeric combinations form high-confidence interactions, in both ROS and No ROS conditions (Fig.1). This is lower than the percentage identified in a similar screen for the family of LRR-RKs (26,4%)^16,48,96^. However, a significantly larger family was tested in that instance, resulting in a dataset in which the cutoff differed considerably from that used in the CRK interactome screen. Moreover, the LRR-RK ECDs interactome screen includes many more well-known interacting pairs, such as ligand-sensing receptors, co-receptors, and regulatory proteins like FLS2-BAK1 and (BAK1-interacting receptor-like kinase 2) BIR2^97–99^. This suggests that the CRK family is not a fully promiscuous receptor set, but rather is organised into preferred dimer and community configurations. The observation that most HCIs are heterodimers is consistent with the previously described dynamics of LRR-RKs^16^. In the heterodimerization model, CRKs may integrate receptor “self-sensing” with cross-family combinatorial signaling. In such a system, homodimers could facilitate baseline or self-referential sensing, whereas heterodimers might broaden ligand recognition or contextual response specificity^100,101^.

Most importantly, the presence of H₂O₂ quantitatively and structurally reshapes the CRK interaction networks (Fig. 1). Although the absolute proportion of HCIs is similar between No ROS and ROS conditions, the presence of ROS changes community structure, connectivity, and centrality, indicating that extracellular redox conditions influence network architecture. Increased inter-community connectivity and redistribution of network hubs under ROS, together with the shift in top interactors from mainly clade IVb/V (no ROS) to clades II/V (ROS), support a model in which ROS acts not only as a simple on/off modulator of individual dimer pairs, but as a global network “re-wiring” factor that promotes the formation of distinct signalling assemblies (Fig.1). For instance, under ROS, previously peripheral nodes such as CRK10 become integrated, and overall connectivity between communities increases. The emergence of CRK17 and CRK29 as dominant hubs with different interacting partners under ROS conditions reinforces this notion of redox-driven shifts in hierarchy. Mechanistically, this assay offers novel insights, as the previously documented ROS-dependent modulation of protein complex formation was predominantly observed in homo-oligomerization, such as in the cases of NPR1 or 2-Cys peroxiredoxin^102–106^. In this study, we have demonstrated differential complex formation events among various CRK members. From a systems perspective, such an architecture would allow flexible redistribution of signalling flow based on redox status, without requiring large-scale changes in receptor abundance. This is also applicable to CRK28, which, in the absence of ROS, remains a homodimer not connected to the network. Its homodimerization has also been demonstrated at the level of the full-length receptor by Yadeta et al. (2018). However, under ROS conditions, homodimerization becomes more robust, and CRK28 interacts with a broader array of CRKs through heterodimerization with CRK17. Such and many more interactions also depend on the spatiotemporal expression of certain CRKs, and therefore, this criterion can be used to narrow down the list of potential interacting proteins (Fig.2).

### Cysteine oxidation as a mechanistic driver of ROS-dependent CRK dimerization

One of the most postulated concepts of ROS perception relies on the ability of cysteine residues to undergo reversible oxidation-reduction reactions via their thiol groups, leading to oxidative post-translational modifications (oxPTMs)^107^. These reversible oxidative changes on cysteines can act as redox switches, rapidly influencing biological processes by altering protein structure, activity, localisation, and binding properties. Redox regulation affects intracellular processes, including transcription factor regulation, oxidoreductase activity (e.g., thioredoxins and glutaredoxins), and defence signalling involving NPR1^29,104,108,109^. Notably, CRK proteins were also postulated to play a significant role in extracellular ROS perception, featuring multiple cysteine residues in conserved (C-X8-C-X2-C) and non-conserved motifs ^110^.

The DAM–MS analysis provides direct evidence that a subset of CRK ECD cysteines undergo reversible oxidative modification in response to H₂O₂ ^106,111–113^. Although only 10 of 32 tested ECDs exhibited differentially alkylated cysteines, and usually only 2–3 per protein, this is compatible with a scenario in which only a few key, strategically positioned cysteines act as functional redox sensors or switches, while most remain stably engaged in structural disulphide bonds (Fig.2). This selectivity parallels broader literature on redox signalling, in which only a fraction of cysteines in a proteome act as bona fide redox switches, characterized by appropriate pKa, solvent exposure, and local environment^36^. The identification of CRK17, CRK26, CRK28 and CRK39 as CRKs carrying cysteines that become more oxidized upon H₂O₂ treatment aligns well with their positioning in the RIA^CRK^ network and, in the case of CRK17 and CRK28, with their roles as hubs in ROS-enriched contexts.

The combination of DAM–MS with solvent-accessible surface area calculations strengthens the argument that at least some of these cysteines (e.g. CRK17 C30, CRK28 C228) are both chemically and structurally poised for redox sensing. High RSA implies accessibility to extracellular oxidants and reductants, while the observed increase in oxidation upon H₂O₂ treatment links this structural property to functional responsiveness. For CRK28, the presence of vicinal cysteines (C228/C229) that are predicted, but not necessarily constrained, to form disulphide bonds is particularly compelling. Vicinal cysteine motifs function as canonical thiol-based redox switches in multiple protein families, toggling between reduced, disulfide-linked and higher oxidation states, which induce conformational changes that control protein activity, as illustrated for Hsp33 and other redox-regulated proteins ^114–116^.

Our data strongly support the idea that C228/C229 constitute such a redox switch in CRK28, controlling receptor dimerization in response to extracellular ROS. Mechanistically, this fits into the broader paradigm of ROS perception via thiol-based switches at the cell surface. In animals, several receptor tyrosine kinases and ion channels are regulated by extracellular cysteine oxidation^117–119^. In plants, the H₂O₂ sensor HPCA1 is regulated by oxidation of extracellular cysteines, which promotes receptor activation and Ca²⁺ influx^85,120^. CRK28 (and possibly other CRKs, such as CRK17) appears to function similarly: ROS alters the redox state of specific ectodomain cysteines, shifting the equilibrium between inactive monomers, pre-formed dimers, and/or higher-order complexes. This change in higher-order structure is likely transmitted to the cytoplasmic kinase domains, thereby modifying downstream phosphorylation patterns and signalling outputs.

### CRK28 as a redox-dependent hub receptor

The choice of CRK28 for deeper functional dissection is well justified by its network properties (ROS-enhanced homodimerization; ROS-induced heterodimerization with CRK17), its expression pattern (strong induction during senescence and flg22-triggered immunity), and its redox-sensitive cysteines. FRET-FLIM analysis confirms that CRK28 forms homodimers and heterodimers with CRK17 at the level of full-length receptors *in planta* ^40,121^ (Fig.3). Importantly, the CRK28–CRK17 heterodimer was not above threshold in the No ROS *in vitro* screen but was recovered in the agroinfiltration system. Agrobacterium-mediated agroinfiltration in Nicotiana is known to trigger PTI-like defense responses, including NADPH-oxidase–dependent production of ROS in the apoplast^122–124^. This discrepancy strongly suggests that physiological redox contexts can reveal interactions that appear weak or absent under artificially controlled *in vitro* conditions, highlighting both a strength and a limitation of the RIA^CRK^ approach.

The cysteine mutagenesis series in CRK28 provides mechanistic insight into how the ECD cysteines contribute to folding, trafficking and dimerization (Fig.3, Supp. Fig.7 and 8). Disruption of most predicted disulphides (e.g. C89/C99, C102/C127, C205/C214, C217/C242) leads to ER retention or irregular PM localisation, consistent with a requirement for these bonds in structural integrity and proper folding. By contrast, the C228A/C229A double mutant maintains correct PM localisation and Hechtian strands, indicating that these vicinal cysteines are not essential for receptor folding or ER quality control. Yet, FRET-FLIM shows that CRK28^C228A,C229A^ fails to homodimerize, despite unaltered localisation. The observation that C228, but not C229, is detectably oxidized in *N. benthamiana* further refines this model: C228 appears to be the functionally redox-active residue, whereas C229 might contribute structurally or electronically to stabilising C228’s redox transitions.

Taken together, these data strongly support the idea that C228/C229 constitute a redox switch governing CRK28 dimerization. Under oxidising conditions, oxidation of C228 (possibly in a dynamic interplay with C229) may favour or stabilise the CRK28–CRK28 and CRK28–CRK17 interfaces; loss of these cysteines abolishes dimerization without gross misfolding. This is a hallmark of regulatory, rather than structural, disulphides and is consistent with the broader hypothesis that CRKs sense extracellular ROS via specific cysteines in their DUF26 domains.

### CRK28 as a regulator of senescence and an autoimmunity-like state

At the whole-plant level, the genetic data indicate that CRK28 functions as a dosage-sensitive regulator of growth, senescence, and immune homeostasis^40,125^. The *crk28* knockout shows moderately enhanced primary root growth and delayed juvenile-to-adult leaf phase transition, together with delayed senescence in the leaf-detachment assay (Fig.4, 5). These phenotypes are consistent with CRK28 normally contributing to growth restraint and the timely initiation of senescence-associated programmes ^40,125,126^.

By contrast, the npro:CRK28-FLAG line, which at 5 weeks exhibits approximately 15-fold higher CRK28 transcript levels than Col-0, displays a striking dwarf, precociously senescing, and necrotic phenotype reminiscent of constitutive autoimmunity^127–129^. Proteomic profiling across development underscores that these morphological changes coincide with a progressive reconfiguration of the proteome (Fig.4, Supp.Fig. 13). In the *crk28* line, developmental reprogramming mainly affects protein synthesis and secondary metabolism. In npro:CRK28-FLAG, late stages of development are dominated by the accumulation of proteins associated with defence, systemic acquired resistance, senescence, and redox homeostasis, along with a coordinated loss of photosynthetic and chloroplast-associated proteins. The concurrent overaccumulation of CRK28 within the same cluster as stress response and senescence-related proteins indicates that elevated CRK28 is not merely a marker but also a driver of this reprogrammed state.

Classical senescence (SAG12, SAG13, LEA, DIN11) are markedly upregulated in npro:CRK28-FLAG leaves at 5 weeks, and chlorophyll measurements confirm accelerated chlorophyll loss (Fig.5). Early and extensive cell death, reduced flg22-triggered ROS burst, and the accumulation of pathogenesis-related proteins and SA-biosynthetic enzymes are all features shared with known autoimmunity mutants, particularly those with misregulated RK or RLP complexes^64,130,131^. Thus, the phenotype of npro:CRK28-FLAG is most parsimoniously interpreted as a CRK28 dosage-dependent autoimmunity-like state in which persistent or excessive CRK28 signalling locks the plant into a high-ROS, defence-primed, senescence-prone mode (Fig.5).

The observation that flg22-triggered ROS production is reduced in npro:CRK28-FLAG, while elevated in *crk28*, is particularly informative. It implies that CRK28 positively constrains canonical early immune ROS bursts, possibly via feedback regulation of NADPH oxidases or apoplastic peroxidases^62,63^. When CRK28 is overabundant at a developmental stage already associated with elevated endogenous ROS (late rosette), this feedback may become maladaptive, shifting ROS production towards sustained rather than transient dynamics and diverting redox resources into chronic defence and senescence pathways. The strong enrichment of peroxidases and glutathione transferases in the npro:CRK28-FLAG proteome is consistent with chronic ROS stress that plants attempt to buffer but fail to fully control. Moreover, CRK28 overaccumulation causes a massive, system-wide rewiring of the plant phosphoproteome, especially at the plasma membrane. This leads to broad activation and reconfiguration of immune, ROS, hormone, and trafficking signalling networks that match the autoimmune-like, early-senescence phenotype.

### CRK28-containing complexes as nodes in stress and membrane trafficking networks

The CRK28 interactome in 5-week-old npro:CRK28-FLAG plants supports a model in which CRK28 functions at the convergence of redox, defence, and vesicle trafficking pathways (Fig.6). Its co-purification with multiple PR proteins (PDF1.2b, PR-5, LTP5) and antioxidant enzymes (SOD1, GSTs, peroxidase 67) suggests that CRK28 resides in, and perhaps helps to organise, an extracellular microenvironment specialised for pathogen defence and ROS detoxification. The association with WAK1 further embeds CRK28 into the cell wall integrity and damage-sensing network, while the interaction with SOBIR1 connects it to RLP-mediated immune complexes and the BIR1–SOBIR1 checkpoint that guards against runaway immunity. Furthermore, both RLK members were identified as hyperphosphorylated at this stage in the npro:CRK28-FLAG line.

Notably, the identification of SNAREs (SYP121, SYP122) and VAMP722 as interacting partners, along with the hyperphosphorylation of these trafficking proteins in npro:CRK28-FLAG plants, suggests a scenario in which CRK28 not only responds to ROS and defence signals but also actively regulates exocytosis and plasma membrane composition (Fig.6). SYP121/SYP122–VAMP722 complexes are integral to the controlled secretion of defense-related cargo to the apoplast ^75,76,132^. Hyperactivation or misregulation of a CRK28–SNARE signalling axis could thus explain the high accumulation of PR proteins at the cell surface and the increased phosphorylation level of trafficking components, aligning with excessive or mistimed vesicle fusion events.

In this integrated view, CRK28 functions as a ROS-tunable hub receptor that (i) modulates its dimerisation with itself and CRK17 in response to the redox state; (ii) coordinates the assembly of PR, antioxidant, and cell wall signalling proteins at the PM–apoplast interface; and (iii) controls the trafficking machinery responsible for delivering these components. Under normal physiological conditions, such a hub promotes efficient, self-limiting defence and regulated senescence. However, under conditions of overexpression and increased endogenous ROS levels, possibly due to RBOHD hyperphosphorylation and differential phosphorylation of PIPs, it may instead trigger a self-amplifying loop that results in autoimmunity.

### Limitations and future directions

Several important considerations require attention. Firstly, the RIA^CRK^ assay depends on ectopically expressed ECDs in Drosophila S2 cells and utilizes a single reactive oxygen species (H₂O₂) at a specific concentration. Other extracellular oxidants (e.g., ·OH, O₂⁻) or pH conditions may further influence CRK–CRK interactions. Expanding the applicability of RIA^CRK^ to multiple redox environments could enhance accuracy and reveal the dynamics of redox network mapping. Secondly, the coverage of cysteines in DA–MS analysis is incomplete, limited by proteolytic constraints and the technical difficulties in capturing reversibly oxidised states. Employing more comprehensive proteomic approaches (such as alternative proteases, targeted probes and more sensitive MS instruments) will facilitate more precise identification of redox-sensitive residues. As follows, the precise structural mechanism by which oxidation of C228/C229 and related cysteines alters CRK28 conformation and dimerisation could be addressed by resolving ectodomain structures in different redox states or as it was demonstrated for CARD1 (HPCA1) Also, in the broader context, it may be interesting to ask whether similar redox-switch cysteine motifs underlie ROS-dependent dimerization in other CRKs identified by the RIA^CRK^ and DA–MS methods, and whether CRK-based redox modules are conserved in other plant species, like crops. It will be intriguing to evaluate whether modifying CRK dosage or redox sensitivity could potentially improve stress resilience without eliciting adverse effects autoimmunity. Lastly, the genetic dissection of single vs. double cysteine mutants (e.g. C228A vs. C229A vs. C228A/C229A) in stable Arabidopsis lines would clarify the individual contributions of each cysteine to signalling in planta. Dissecting the functional relationships between CRK28 and its identified interacting partners (WAK1, SOBIR1, SNARE) could reveal how CRK28 integrates cell wall, immune, and trafficking inputs.

## Concluding remarks

This work provides a clearly delineated mechanistic framework that associates ROS-dependent CRK extracellular domain (ECD) dimerization, cysteine redox chemistry, receptor complex assembly, and the regulation of developmental and defence responses. CRK28 is identified as a prototypical redox-sensitive CRK, utilizing specific, solvent-exposed cysteines to detect extracellular ROS and modulate receptor–receptor interactions, thereby balancing growth, senescence, and immunity. The comprehensive RIA^CRK^ map and spatiotemporal networks serve as valuable tools for elucidating how plants interpret complex ROS signals through a modular receptor system, as they offer many more relevant candidates besides CRK28. These resources additionally facilitate the rational engineering of redox-sensing mechanisms to enhance stress resilience while avoiding the induction of autoimmunity.

## Materials & Methods

### Molecular cloning of CRK extracellular domains

The ECDs of CRKs were cloned by Seamless Ligation Cloning Extract (SLiCE)^133^. This is a recombination system in which the vector backbones of pECIA 2 and 14 (Addgene #47032 and #47051, respectively) are amplified with a reverse primer on the existing BiP signal sequence and a forward primer on the C-terminal epitope tags specific to each vector. Primers were designed to have a sequence partially homologous to the desired boundaries of each CRK ECD amplified from a cDNA library of *Arabidopsis thaliana* ecotype Col-0 with additional 15-20 bp of flanking sequences with homology to the vector’s ends (Supp. Table 1). For CRK11 and CRK16, synthetic gene fragments were assembled with homologous flanking sequences to pECIA2 and pECIA14 using NEBuilder^®^ HiFi DNA Assembly kit (New England Biolab) according to the manufacturer’s specifications. The boundaries of CRK ECDs were determined by using bioinformatic tools^134^. The plant signal peptide, which usually comprises approximately the first 20 amino acids, was removed, and ECDs were amplified until just before the TM. These boundaries were confirmed by visual inspection of the amino acid sequence. Sanger sequencing was used to confirm the presence and integrity of each insert. For the cysteine redox state determination using the Differential Alkylation Method (DAM), the same CRK ECD sequences were cloned into the recombinant pECIA2 vector with only a minimal V5 tag. Amplification was done using Phusion Flash Mastermix (Thermo Fisher Scientific) according to the manufacturer’s instructions for 2-step PCR.

### Secreted expression of CRK extracellular domains

The ECDs were cloned into pECIA2 and pECIA14 vectors for expression as baits and preys, respectively. These vectors were expressed according to the procedure described in Smakowska-Luzan, 2018, using transient transfection of Drosophila S2 cells (94-005F, Expression Systems) cultured at 27°C in Erlenmeyer flasks with baffles and constant mild shaking. Transfection was done using the Effectene kit (Qiagen) according to manufacturer’s specifications. During transfection, the temperature was lowered to 21°C and the incubation was done without shaking. Twenty-four hours after transfection, protein expression was induced with 1 mM CuSO_4_. Three days after induction, the culture was spun down to precipitate the cells and the supernatant was collected. In order to prevent protein degradation, 0.02% NaN_3_ and cOmplete™ EDTA-free Protease Inhibitor Cocktail (Roche) was added to the medium (ESF 921, Expression Systems) containing the recombinant ECDs and stored at 4 °C before use. Protein expression was assessed by western blotting using anti-V5 antibodies (Invitrogen) for the baits or by alkaline phosphatase activity quantification using the KPL BluePhos® Microwell Phosphatase Substrate System (SeraCare) for the preys.

### RIA^CRK^ screen

The wells of Protein-A-coated plates (Thermo Fisher Scientific) were washed with 100 µL of TBS + 0.1% Tween-20 for 10 minutes with mild shaking before use. 25 µL of bait proteins were diluted with 75 µL of TBS supplied with 0.5 mM of dithiothreitol (DTT) and captured directly on each well of 96-well protein-A-coated plates by incubating overnight (ON) at 4°C with mild shaking. Parallelly, the prey proteins that were going to be used the next day were also supplied with 0.5 mM of DTT and incubated ON at 4°C. After incubation, bait proteins were discarded, and 25 µL the prey proteins fused to the alkaline phosphatase were added to the bait-coated wells, diluted with 75 µL of TBS and incubated for 2h at room temperature (RT) with mild shaking. During prey incubation, the effect of H_2_O_2_ on the protein interactions was tested by adding either nothing (No ROS) or 100 µM H_2_O_2_ (ROS) to the reaction media. After incubation, the media with the prey proteins was carefully removed before adding 100µL KPL BluePhos® Microwell Phosphatase Substrate System (SeraCare). The plates were then incubated for 2h at RT, and alkaline phosphatase activity was recorded by measuring the absorbance at 650 nm using a Synergy H4 Multi-Mode plate reader (BioTek). Images of the 96-well plates were also acquired for visual quality control. The complete set of raw absorbance values was combined into a matrix dataset and then subjected to post-experimental statistical analysis to remove both false positive and false negative interactions.

### Modelling with AlphaFold multimer

All AlphaFold predictions of CRKs ECDs were done using AF v2.2.2, parameters were set to default. For monomer and multimer predictions, sequences of CRK ECDs cropped to the structural part of the DUF26 domains as input. For the monomer and multimer 5 models were generated each. The ipTM scores of pairs of CRKs used for multimer predictions were plotted as a heatmap in GraphPad Prism v10.2.3.

### RIA^CRK^ Interactome Data Analysis

All absorbance values for each testing interaction were combined into a data matrix. A two-way median polish was used to eliminate bias due to differential background binding capacities and make plates comparable ^135,136^. The residuals were then used to calculate the median and median absolute deviation (MAD) for the distribution. The MAD was used for the calculation of modified *Z*-scores for each individual interaction measured. To ensure that only bidirectional interactions were used, we calculated the geometric mean modified *Z*-score of the interaction as measured in the bait–prey and prey–bait orientations. Any value for which the geometric mean product of the *Z*-scores was greater than 1 was considered significant for the purposes of network construction.

### Network construction and analysis

The network was constructed using the igraph package (http://igraph.org/r/) in the R programming environment (https://www.r-project.org/). Clusters of interacting proteins in the network were identified using the WalkTrap algorithm as implemented in igraph using a random walk length of 8^137^. The PageRank algorithm was used as a centrality measure within the network, as implemented in igraph, with the node size set to be relative to the PageRank score ^121^. Betweenness centrality was calculated as implemented in igraph.

### Construction of developmental and flg-22 triggered interaction maps

The RIA^CRK^ networks were filtered based on the expression of each gene in two different developmental stages (seedlings and senescence) and after 90 minutes of treatment with flg22. Expression of 40 CRKs was retrieved from TravaDB ^121^ and ^57^using the default median-ratio method, and then the Raw normalisation settings that provide the number of Absolute counts for each gene and each tissue were assembled as a matrix. The average of counts was calculated for all 40 genes in all tissues (195), and this number was used as a cutoff for considering a gene expressed. For each developmental subnetwork, the interactors of HCIs in RIA^CRK^ were evaluated as expressed or not expressed if the absolute expression value for that interactor at that developmental stage was above or below the cutoff. An interaction was considered valid and plotted in the subnetwork only if both interactors of the HCI were present in that developmental stage. The subnetworks were constructed using the igraph package in R as described above.

### Cysteine redox state determination using the Differential Alkylation Method (DAM)

The redox state of the different cysteines was determined using the differential alkylation method which was executed on top of (ethanol washed Pall 3K omega) 10 kDa membranes to make separation of reagents and proteins possible by centrifugation^138^. After each incubation step, the reaction mixture was centrifuged at 4 °C by 12.000 *g through the filter in 30 min or longer when necessary to make all liquid pass.

All samples were prepared in triplicate. One hundred ul of cleared insect cells medium containing 10 ug of proteins, as determined from a BSA Bradford assay, were treated with buffer only as a control or buffer with 100 mM H2O2 for 2 hours at room temperature^139^. Acrylamide was added up to a concentration of 50 mM as the first alkylation agent. After gently mixing at room temperature for 10 min, the whole sample was transferred to an ethanol washed Pall 3K omega filter fitted into a 1.5 ml microcentrifuge tube which was centrifuged. One hundred ul 50 mM ditiothreitol reductant in 100 mM Tris pH8 was added, incubated at room temperature for 30 min and centrifuged. Proteins were unfolded by adding 200 ul 8M urea in 100 mM Tris pH8, gently shaken and centrifuged until dry. One hundred ul 50 mM Iodoacetamide in 100 mM Tris pH8 (as a second alkylation reagent) was added, incubated at room temperature for 10 min and centrifuged until dry. The filters were washed by adding 200 ul 50 mM ABC and centrifuged. The filter cups were transferred to clean 2 ml low binding Eppendorf microcentrifuge tubes and 100 ul 5 ng/ul bovine sequencing grade trypsin (Roche 11 047 841 001) in ABC were added and, while gently shaking, incubated at room temperature for at least 15 hours. Any liquid that had passed through the filter was pipetted back on top of the filter and peptides were de-glycosylated by adding 2 µl Glyco3 buffer and 5 units PNGaseA (New England Biolabs). Samples were gently shaking at 37 °C and incubated for 1 hour. Three ul 10% TFA was mixed into the solution on top of the filter and acidified peptides were centrifuged through the filter. The filtrate was dried in a vacuum concentrator, re-dissolved in 10 ul 1 ml/l HCOOH in water, frozen and kept at -20 °C until samples were measured by nLCMS.

Five ul of peptide samples were loaded directly onto a 0.10 * 250 mm Reprosil Saphir 100 C18 1.5 µm beads analytical column (prepared in-house) at a constant pressure of 1050 bar (flow rate of circa 700 nL/min) with 1 ml/l HCOOH in water and eluted at a flow of 0.5 ul/min with a 50 min gradient from 0 to 29% acetonitril in water with 1 ml/l formic acid using a Thermo Vanquish Neo nanoLC. An electrospray potential of 3.0 kV was applied directly to the eluent via a stainless steel union connecting the nLC outlet tubing and the analytical column. Full scan positive mode FTMS spectra were measured between m/z 380 and 1400 on a Exploris 480 (Thermo Fisher Scientific, Bremen, Germany) in the Orbitrap at resolution 60000 using a DDA method. MS and MSMS AGC targets were set to 300%, 100% respectively or maximum ion injection times of 50 ms (MS) and 30 ms (MSMS) were used. HCD fragmented (Isolation width 1.2 m/z, 28% normalized collision energy) MSMS scans in a cycle time of 1.1 s of the most abundant 2-5+ charged peaks in the MS scan were recorded in data dependent mode (Resolution 15000, threshold 5e4, 12 s exclusion duration for the selected m/z +/- 10 ppm).

LCMS runs with all MSMS spectra obtained were analysed with MaxQuant 2.0.3.0^140,141^ using the “Specific Trypsin/P” Digestion mode with maximally 2 missed cleavages for the Andromeda search engine (First search 20 ppm peptide tolerance, main search 4.5 ppm tolerance, MSMS fragment match tolerance of 20 ppm, Carbamidomethyl/Propionamide (C) set as possible variable modification, while variable modifications were also set for Protein N-terminal Acetylation and M oxidation which were completed by non-default settings for de-amidation of N and Q, the maximum number of modifications per peptide was 5 ^140,141^. A Drosophila melanogaster (UP000000803) and a Arabidopsis thaliana Cysteine Rich Kinases (containing 32 sequences and including a his tag as they were produced in the insect cells) databases downloaded from Uniprot (http://www.uniprot.org) were used together with a contaminants database that contains sequences of common contaminants like Trypsins (P00760, bovin and P00761, porcin) and human keratins (Keratin K22E (P35908), Keratin K1C9 (P35527), Keratin K2C1 (P04264) and Keratin K1CI (P35527)). The “label-free quantification” as well as the “match between runs” options were enabled. De-amidated peptides were allowed to be used for protein quantification and all other quantification settings were kept default.

The nLC-MSMS system and data quality was checked with PTXQC ^142^ using the MaxQuant result files^142^.

Result filtering and further bioinformatic analysis of the MaxQuant/Andromeda workflow output and the analysis of the abundances of the identified proteins were performed with the Perseus 1.6.2.1 module^143^. Accepted were peptides and proteins with a false discovery rate (FDR) of less than 1% and proteins with at least 2 identified peptides of which at least one should be unique and at least one should be unmodified. Reversed hits were deleted from the MaxQuant result tables.

For the calculation of percentages of reduced cysteines (alkylated with acrylamide), the ‘evidence.txt’ file was used. Intensities of Reduced cysteines (Acrylamide modified cysteines) divided by intensities of Reduced + Oxidized cysteines (Acrylamide + IodoAcetamide modified cysteines) * 100% were calculated per peptide per condition while summing up intensities found for different charge states. The mass spectrometry proteomics data have been deposited to the ProteomeXchange Consortium via the PRIDE partner repository with the dataset identifier PXD075564.

### Cloning and baculovirus stock production

For the production of baculovirus stocks, the coding DNA for the extracellular domain of CRK17 was amplified with an N-terminal melittin signalling peptide and a C-terminal 9xHIS tag. The construct was first inserted into pDONR221 and subsequently pOET1 transfer vector (200101, Oxford Expression Technologies) using Gateway cloning according to the manufacturer’s instructions (Invitrogen). Baculovirus stock was produced in *Spodoptera frugiperda* Sf9 insect cells (Expression Systems) using the FlashBac Ultra system (Oxford Expression Technologies) according to the manufacturer’s instructions. Viral titres were determined using Sf9-EasyTiter cells ^144^.

### Protein expression and purification

Protein was expressed in *Trichoplusia ni* (TniH5) cells (Invitrogen) infected with baculovirus. Cells were cultured for 72 h at 27 °C. Cells were removed from the protein-containing supernatant by centrifugation (1.500 g, 30 min). The supernatant was subjected to affinity chromatography with Ni Sepharose excel resin (Cytiva Life Sciences). The resin was washed with wash buffer (20 mM Tris, 500 mM NaCl, 10 mM Imidazole, 1 mM DTT). Subsequently, the protein was eluted from the column (Elution buffer, 20 mM Tris, 500 mM NaCl, 350 mM imidazole, 1 mM DTT). Protein was further purified on a preparative grade Superdex S200 column in HBS (20 mM HEPES, 150 mM NaCl).

### Thermal unfolding assay

To assess the effect of DTT and H_2_O_2_ treatments on ECD stability, nanoDSF assays were performed on purified CRK-ECDs. Protein solutions at a concentration of 0.4 mg/mL in TBS, MgCl, 0.1% pluronic acid were measured in capillaries in triplicate using a Prometheus Panta instrument (NanoTemper). Measurements were conducted over a 20 to 95 °C temperature range at a ramp rate of 0.5 °C/min. Results were processed in Panta analysis software to determine the T_m_ from the 350nm/330nm ratio and plotted in GraphPad Prism.

### Transient expression in Nicotiana benthamiana and localization experiments

CRK28 genomic DNA and xx (ER-marker) were cloned into modified pGIIB expression vectors. The modified vectors had a 3xFLAG-mScarlet-I and mNeonGreen-V5 tag respectively^145^.

CRK28 cysteine-to-alanine substitution mutations and the K377N mutation were introduced using site-directed mutagenesis or by primer amplification incorporating the point mutation, followed by vector reassembly using HiFi (NEB).

CRK28 constructs were transformed into A. tumefaciens strain GV3101-pSoup. Infiltration solution was prepared in 10 mM MES, 10 mM MgCl, 100 uM acetosyringone, Agrobacterium CRK constructs to an OD of 0.3, and 0.5 for P19. Localization of CRK28 and cysteine mutants was measured using a Leica SP8 confocoal microscope. mScI was excited at 561 nm and emission was detected using hybrid detectors (580-630 nm). mNG was excited at 488nm, and emission was measured 490-530 nm. A 63x water NA 1.2 objective was used. For each sample/mutant leaf discs of four-week-old N.benthamiana plants 2 days DPI, three images per plant, three different plants.

### FRET-FLIM in Nicotiana benthamiana

CRK28, CRK28^C228AC229A^ and CRK17 cDNA from Arabidopsis were cloned into estradiol-inducible expression vectors described in ^146^. Transient expression of CRK17 and CRK28 variants in N. benthamiana was performed as described in ^147^. Lifetime measurements were performed using a Zeiss LSM 780 confocal microscope (40× water immersion objective, Zeiss C-PlanApo, NA 1.2). For Time-Corelated-Single-Photon-Counting (TCSPC) measurements a PicoQuant Hydra Harp 400 (PicoQuant, Berlin, Germany) was used. “Photon counting was performed with a picosecond resolution. GFP was excited with a 485 nm (LDH-D-C-485, 32 MHz, PicoQuant, Berlin, Germany) pulsed polarized laser. The laser power at the objective lens was adjusted to 1 µW. mCherry was excited with a 561 nm laser at 1% power. Light, emitted from the sample, was separated by a polarizing beam splitter and a LP610 beam splitter before photons were selected with a fluorophore-specific band-pass filter. Photons were detected in both donor and acceptor channel simultaneously with Tau-SPADs (PicoQuant, Berlin, Germany). Images were acquired at zoom 8 resolution of 256x256pixel with a pixel size of 0.1 µm and a pixel dwell time of 12.54 µs and laser repetition rate of 32 MHz. Photons were collected over 40 frames. Before image acquisition, the system was calibrated. First, the objective was adjusted to reach a maximal count rate. Fluorescence-Correlation-Spectroscopy (FCS) curves of Rhodamine110 dye and water were acquired to monitor the system function. Internal Response Function (IRF) for each laser were determined by measuring the fluorescence decay of quenched erythrosine in saturated potassium iodide using the same hardware settings as for the FRET pair.

In order to determine the average lifetime of recombinant proteins, regions of interest (ROIs) of the plasma membrane in the FLIM image were selected manually. The fluorescence decay times of the ROIs were analysed using SymPhoTime FLIM analysis software (SymPhoTime 64, version 2.4; PicoQuant, Berlin, Germany). Time-correlated single photon counting (TCSPC) bins of donor channel 1 (parallel light) and channel2 (perpendicular light) were binned 16 times. Pixels above the pile-up limit (10% of the lase repetition rate) and chloroplasts were removed manually. The data was fitted using the grouped FLIM image analysis tool (n-exponential reconvolution, n=2). The intensity-weighted lifetime was considered as the sample’s apparent lifetime. All samples were measured in three independent experiments.

### Plant growth and conditions

Arabidopsis plants were grown on soil at 21°C under long-day (16 h light: 8 h dark) conditions and 60% humidity for 3, 4, and 5 weeks. Rosettes that displayed a representative phenotype were selected for whole-rosette images and leaf panels. 8-12 plants per genotype were used for detached leaf senescence measurements. For seedling experiments, seeds were sterilized and sown on 1/2 Murashige-Skoog media. Seedlings were grown for 2 weeks at 21 °C under long-day (16 h light: 8 h dark) conditions in sealed plates. T-DNA insertion line of CRK28 (*crk28*, SALK_085178) in the Col-0 background was obtained from the Arabidopsis Biological Resource Centre. For overexpression of CRK genes, we used a stable Arabidopsis line harbouring the native CRK28 promoter driving expression of CRK28 transcriptionally fused to a FLAG tag in the *crk28* T-DNA background (*npro*:CRK28-FLAG), as published by ^40^.

### Extraction of mRNA and qRT-PCR

RNA was extracted from rosette leaves using the RNeasy Plant Mini Kit (Qiagen) according to the manufacturer’s instructions. DNase treatment was performed using the RNase-Free DNase Set (Qiagen) prior to cDNA synthesis, which was achieved with the iScript Select cDNA Synthesis Kit (Bio-Rad). Finally, relative expression levels were determined by quantitative reverse transcription polymerase chain reaction (qRT-PCR) using iQ SYBR Green Supermix with the CFX384 Real-Time System (Bio-Rad). *Ribosomal protein S26C* (*RPS26C*) and *20S PROTEASOME ALPHA SUBUNIT C1* (*PAC1*) were used as reference genes for normalization. The primers used for RT-qPCR are listed in Supp. Table 1.

### Leaf detachment assay

For detached leaf senescence measurements, the 3rd and 4th leaves were harvested from 5-week-old plants. Leaves were placed on top of one layer of Whatman filter paper soaked with ½ MS pH 5.8 and sealed inside of a petri dish with parafilm. Pictures were taken daily to follow senescence symptoms (Z. Zhang & Guo, 2018). Representative pictures were selected to assemble the figures. For semi-quantitative measurements of senescence progression, the green-to-red pixel ratio of the leaves in the images was quantified in ImageJ using the GreenLeafVI plugin from Leiden University^148,149^.

### Cell death assay in Arabidopsis leaves

Cell death was assessed by an increase in water conductivity due to ion leakage of Arabidopsis leaves. Assays were performed according to Lapin et al. (2019) with minor modifications. 2-3 leaf disks of 4mm diameter were harvested per leaf from the 5th and 6th leaves from 5-week-old plants for a total of 24 leaf disks per sample. Leaf disks were placed individually in 100 µL of MQ water in 96-well plates and incubated by shaking at RT ON. After incubation, the water was replaced with 100 µL of fresh MQ and the water conductivity was measured at times 0h and 6h using a LAQUAtwin EC-11 conductometer (Horiba).

### ROS burst assay

ROS burst assay was performed according to ^149^. 2 leaf disks of 4mm diameter were harvested from the 7th and 8th leaves from 5-week-old plants for a total of 24 leaf disks per sample. Leaf disks were placed individually in 100 µL of MQ water in 96-well plates and equilibrated by shaking at RT ON in the dark. After incubation, the water was replaced by 100 µl of luminescence measurement solution (20 µM luminol L-012 (Fujifilm); 1 µg/ml horseradish peroxidase (MERCK) +/- 1 µM flg22 (GenScript) and luminescence was immediately measured for 1 h using a Synergy H4 Multi-Mode plate reader (BioTek).

### Chlorophyll content measurements

Chlorophyll measurements were performed according to ^150^. Leaf disks from ion leakage and ROS burst experiments were incubated with 96% ethanol ON at RT in the dark on a shaking platform. Afterwards, we measured absorbance at 665nm and 649nm, and chlorophyll was calculated with the following formula:

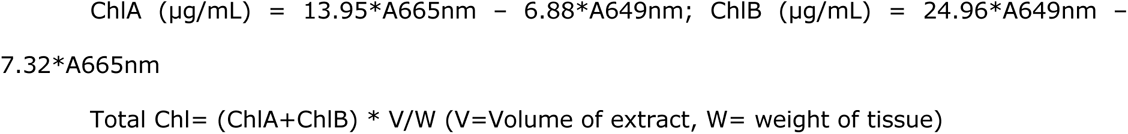

### Protein extraction and Affinity Purification (AP) from Arabidopsis

For IP-MS from mature tissue, 3 whole rosettes from 3-, 4-, and 5-week-old plants were flash-frozen in liquid N_2_ for one technical replicate. Each sample consisted of 3-4 technical repeats. Frozen tissue was ground to a fine powder in liquid N_2_ and extraction buffer (50 mM Tris-HCl, pH 7.5, 150 mM NaCl, 10% glycerol, 10 mM DTT, 10 mM EDTA, 1 mM NaF, 1 mM Na_2_MoO_4_, 1% (v/v) cOmplete Tablets EDTA-free (Sigma-Aldrich), and 1% (v/v) IGEPAL CA-630 (Sigma-Aldrich)) was added at 1 ml per ml of tissue powder. Samples were incubated for 1h at 4 °C with constant rotation. Samples were then centrifuged for 15 min at 16,000 g at 4°C. The supernatant was diluted 1:1 with washing buffer containing 50 mM Tris-HCl, pH 7.5, 150 mM NaCl, 1% PMSF, and 0.1% IPEGAL. The diluted supernatants were then incubated for 2h at 4 °C with 40 µL anti-FLAG agarose beads slurry (DYKDDDDK Fab-Trap® Chromotek) with constant rotation. Following incubation, beads were sedimented by centrifugation for 2 min at 1000g at 4°C and then transferred to 1.5 ml microcentrifuge tubes. Beads were then washed twice with 1 mL of washing buffer and twice with 1 mL of washing buffer without IPEGAL before mass spectrometry (MS) sample preparation.

### MS sample preparation with on-bead digestion of the AP samples

Beads from IP samples were washed twice with 1 mL of 50 mM ammonium bicarbonate (ABC) and resuspended in 90 µL of 50 mM ABC. 10 µL of 150 mM DTT was added, and the mixture was incubated for 30 minutes at 45 °C. After incubation, 10 µL of 200 mM iodoacetamide (IAA) was added and incubated for 30 minutes at room temperature (RT). 10 µL of 200 mM L-cysteine (Biochemika) and 10 µL of 10x diluted in ABC sequence-grade trypsin (0.05 µg/µL) were added, mixed and incubated overnight (ON) at RT with mild shaking. The trypsin digestion was stopped by adding 10% trifluoroacetic acid (TFA) to decrease the pH to 3. The supernatant was transferred to a low-bind microcentrifuge tube, carefully avoiding any beads, centrifuged for 10 minutes at 1000g, and then transferred again to a new low-bind tube to avoid any beads in the sample.

### MS sample preparation for shotgun proteomics

Proteomics samples were prepared following the filter-assisted sample preparation protocol. (FASP)^151^. First, samples were reduced by adding 15 mM dithiothreitol (DTT) and incubated for 30 minutes at 45 °C. Then, samples were alkylated by 20 mM acrylamide and incubated for 30 minutes at RT. Then, samples were digested with sequencing-grade trypsin (0.05 µg/µL) at RT.

### Protein extraction and MS sample preparation for phosphoproteomics

For phosphoproteomics, samples were prepared and analysed according to ^152^. For phosphopeptide enrichment, ground Arabidopsis rosettes were suspended in an extraction buffer with 100 mM Tris-HCl pH 8.0, 7 M Urea, 1% Triton-X, 10 mM DTT, 10 U/ml DNase I (Roche), 1 mM MgCl_2_ and1% benzonase (Novagen). The suspended lysate was sonicated using a cooled (4°C) water bath sonicator (Qsonica) using 30 cycles of 30 seconds on and 30 seconds off at 90% amplitude. Lysate was subsequently spun down using a cooled (4°C) tabletop centrifuge at 20.000 g for 30 min. After centrifugation, supernatant was collected and an extra 1% (v:v) of benzonase was added and incubated for 30 min at RT. Acrylamide was added to 50 mM and incubated for an extra 30 minutes at room temperature. After alkylation, proteins were precipitated using methanol/chloroform. To the lysate, 4 volumes of methanol, 1 volume of chloroform and 3 volumes of MQ water was added with rigorous vortexing in between. Lysate was centrifuged for 10 min at 5000 g. After centrifugation, the top layer was discarded, and 3 volumes of methanol were added to further precipitate the protein layer by centrifugation for 10 min at 5000 g. After centrifugation, the supernatant was discarded, and protein pellet was air dried. Proteins were next resuspended in 50 mM ABC and sonicated at 4°C in a water bath sonicator (Qsonica) using 30 cycles of 30 seconds on and 30 seconds off at 90% amplitude. After sonication, protein concentration was measured by Bradford reagent (Biorad). For every biological replicate, 500 mg protein was digested with sequencing grade trypsin (1:100 trypsin:protein; Roche) ON at RT. Next, peptides were desalted and concentrated using C18 microcolumns. For peptide desalting and concentrating, disposable 1000 ml pipette tips were fitted with 4 plugs of C18 octadecyl 47 mm Disks 2215 (Empore™) material and 1 mg:10 mg of LiChroprep® RP-18 (Merck):peptides. Tips were sequentially washed with 100 % methanol, 80 % Acetonitrile (ACN) in 0.1% formic acid and twice equilibrated with 0.1% formic acid. All chromatographic steps were performed by centrifugation for 4 min at 1500 g. After equilibration, peptides were loaded for 20 min at 400 g. Bound peptides were washed with 0.1% formic acid and eluted with 80% ACN in 0.1% formic acid for 4 min at 1500 g. Eluted peptides were suspended in loading buffer (80 % ACN, 5% TFA). For phosphopeptide enrichment, MagReSyn® Ti-IMAC beads (Resyn bioscience) were used. For every reaction, a 1:4 peptide:bead ratio was used. Beads were equilibrated in loading buffer, resuspended peptides were added and incubated for 20 min at RT with slow mixing. After 20 min, bead-bound phospho- peptides were washed once in loading buffer, once in 80 % ACN, 1 % TFA, and once in 10 % ACN, 0.2 % TFA. After washing, phosphopeptides were eluted twice with 50 µl 1 % NH_4_OH. After the last elution, phosphopeptides were acidified using 10 % formic acid. Phosphopeptides were subsequently concentrated using C18 microcolumns. For peptide desalting and concentrating, disposable 200 ml pipette tips were fitted with 2 plugs of C18 octadecyl 47 mm Disks 2215 (Empore™) material and 1 mg:10 mg of LiChroprep® RP-18 (Merck): peptides. Tips were sequentially washed and equilibrated as described above. After equilibration, peptides were loaded for 20 min at 400 g. Bound peptides were washed with 0.1%formic acid and eluted with 80 %ACN in 0.1 % formic acid for 4 min at 1500 g. Eluted peptides were subsequently concentrated using a vacuum concentrator for 30-60 min at 45°C and resuspended in 15ml of 0.1 % formic acid. Phosphopeptides were further prepared for MS according to the filter-assisted sample preparation protocol (FASP)^152^.

### MS measurements

Mass spectrometry on AP-MSMS and phosphoproteomics samples were carried out on an Exploris 480 (Thermo Scientific) coupled to a Vanquish Neo (Thermo Scientific) as previously described^153^. For shotgun proteomics measurements 1.5 µl samples were loaded on a Pepsep column 25 cm x 75 µm x 1,5 µm (Bruker) at a constant pressure of 800 bar with 1ml/l HCOOH in water and eluted at a flow of 0.3µl/min with a 60 min linear gradient from 2% to 35% acetonitrile in water with 1 ml/l HCOOH with a nanoElute2 (Bruker). Data was acquired using an pydiAID optimized diaPASEF acquisition scheme^154^. The collision energy was decreased as a function of the IM from 59 eV at 1/*K*_0_ = 1.6 V cm^−2^ to 20 eV at 1/*K*_0_ = 0.6 V cm^−2^, and the IM dimension was calibrated with three Agilent ESI Tuning Mix ions (*m/z*, 1/*K*_0_: 622.02, 0.98 V cm^−2^, 922.01, 1.19 V cm^−2^, 1221.99, and 1.38 V cm^−2^).

### Raw data processing, quantification and differential expression analysis

Phosphoproteomic acquired data was analyzed with MSFragger version 23 using the built-in “LFQ-phospho” workflow using standard settings and min prob localisation set to 0^155^. AP-MSMS acquired data was analyzed with MSFragger version 23 using the built in “LFQ-MBR” workflow using standard settings^155^. For shotgun proteome acquired data, Bruker .d files were analyzed using DIA-NN version 2.0.2^156^. Spectral libraries were predicted from *Arabidopsis thaliana* (UP000006548) proteome retrieved from UniProt. MSFragger combined_ion.tsv or DIA-NN report.parquet files were used as input for quantification using DirectLFQ with an in house generated Python script^157^. Raw mass spectrometry data is deposited on Pride with accession number: PXD075696. Differential expression analysis was conducted using the DEP/DEP2 packages in R^158^. DirectLFQ calculated intensities were log2 transformed and filtered to have at least 75% valid values in at least one condition. Remaining missing values were imputed using a mixed imputation approach were values missing at random (MAR) were imputed with knn and values missing not at random (MNAR) were imputed based on quantile regression. Differentially expressed proteins were inferred using limma^152,159^. Clustering of time series protein expression profiles were conducted using the Minardo-model as previously described. Gene ontology enrichment was performed using ShinyGO^159^.

## Acknowledgements

We would like to thank Virendrasinh Khandare for his support with mapping cysteine oxidative modifications. We would particularly like to thank Eduard Sabido Aguade and Guadalupe Espadas-Garcia from the UPF Proteomics Unit, Centre for Genomic Regulation (CRG), Barcelona, Spain, for testing the ETHCD fragmentation method to detect oxidative cysteine modifications. We thank Michael Wrzaczek for sharing the transgenic lines created by ^40^.

## Author contributions

S.M.R designed, performed and analyzed the RIA screen, physiological assays and initial proteomics experiments. M.R., R.L., and J.L. performed proteomics experiments and analyzed the data. S.B. conducted a mapping screen for cysteine oxidative modification, and E.S.L. quantified the data. J.S performed AF modelling and CRK28 localization studies. J.M, R.S., and D.H. performed FRET-FLIM studies. J.W provided training and guidance for microscopy-related experiments. C.B. and M.v.O. provided resources and training on large-scale expression of CRKs. W.V. and J.S performed CRK17 large expression, purification and nDSF. A.T.N and G.A.M performed computational approaches. E.S.L supervised the project. S.M.R and E.S.L. wrote the manuscript with input from all authors.

