## Supplementary material for "Redox-dependent extracellular interaction networks of Cysteine-rich Receptor-Like Kinases": Supp. Figures

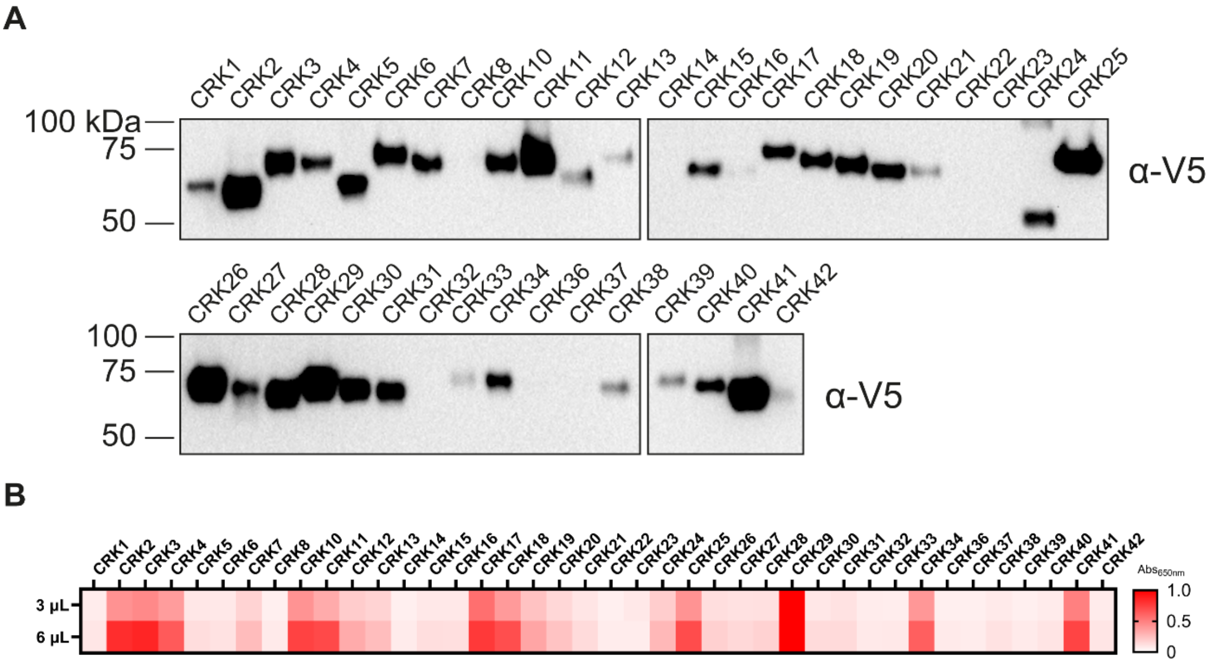


**Supplementary Figure 1. Quality control of bait and prey CRKs used in this study.** A) Western blots of baits detected with anti-V5-HRP antibodies. B) AP activity measured at 650 nm of 3 and 6 µL of prey media into 100 µL of KPL substrate. The scale bar represents absorbance measured at 650 nm.

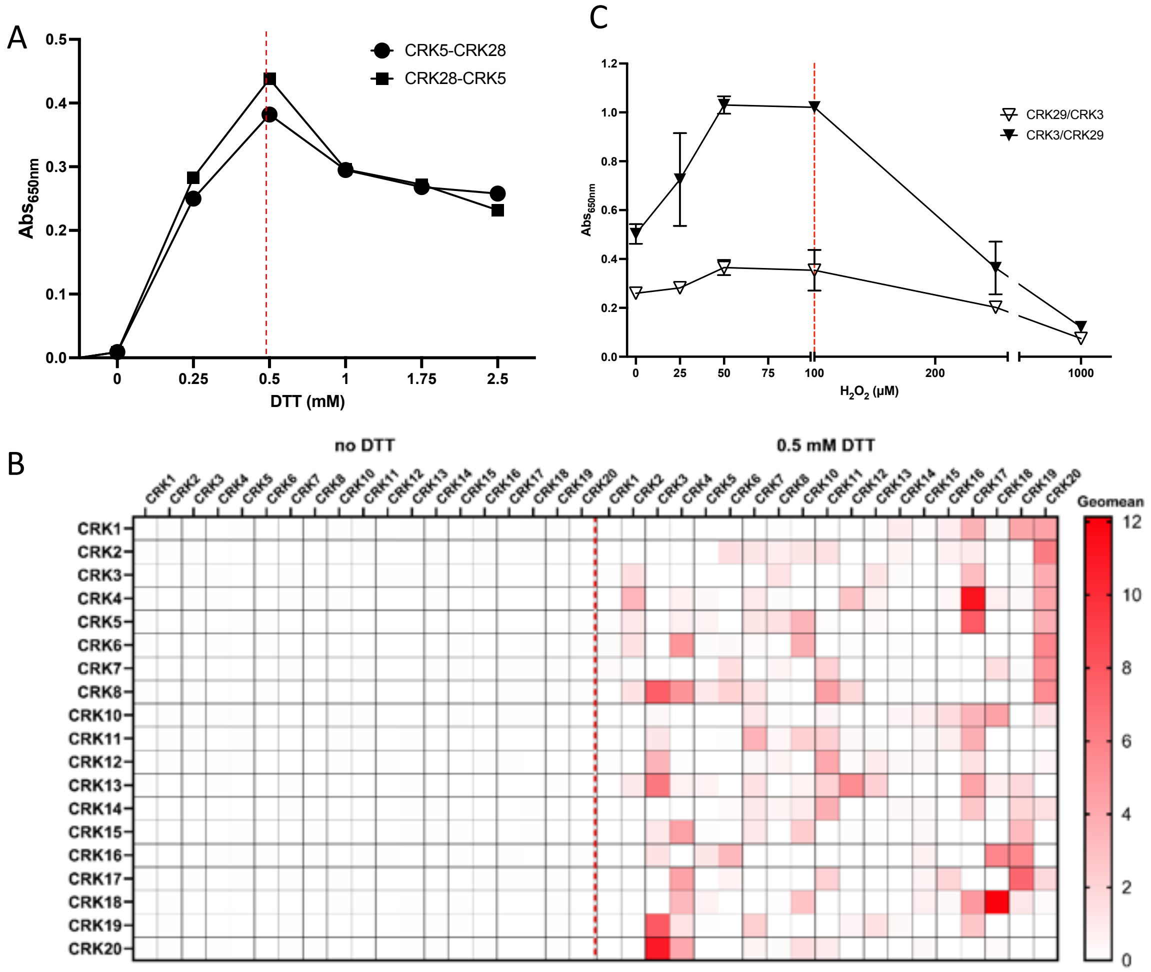


**Supplementary Figure 2**. **Optimizations to the CSI assay to adapt it for CRK ECDs interaction screen with $H_2O_2$ treatment.** A) CSI assay between different CRK bait-prey pairs tested on increasing concentrations of DTT. Each interaction strength is presented after subtracting the signal from the respective prey-alone control with the appropriate concentration of DTT. B) Heatmap of a subset of CRK-CRK interactions in the presence or absence of 0.5 mM DTT. Raw absorbance was subtracted by their respective prey-only signal and normalised by a 2-way median polish to generate modified Z-scores. The scale bar represents the GeoMean of modified *Z*-scores of reciprocal interactions. C) CSI assay between CRK3-CRK29 bait-prey pair tested on increasing concentrations of $H_2O_2$ reciprocally. Each interaction strength is presented after subtracting the signal derived from the respective prey alone with the appropriate concentration of $H_2O_2$. The dashed red line in A) and C) represents the optimal concentration for their respective treatments.

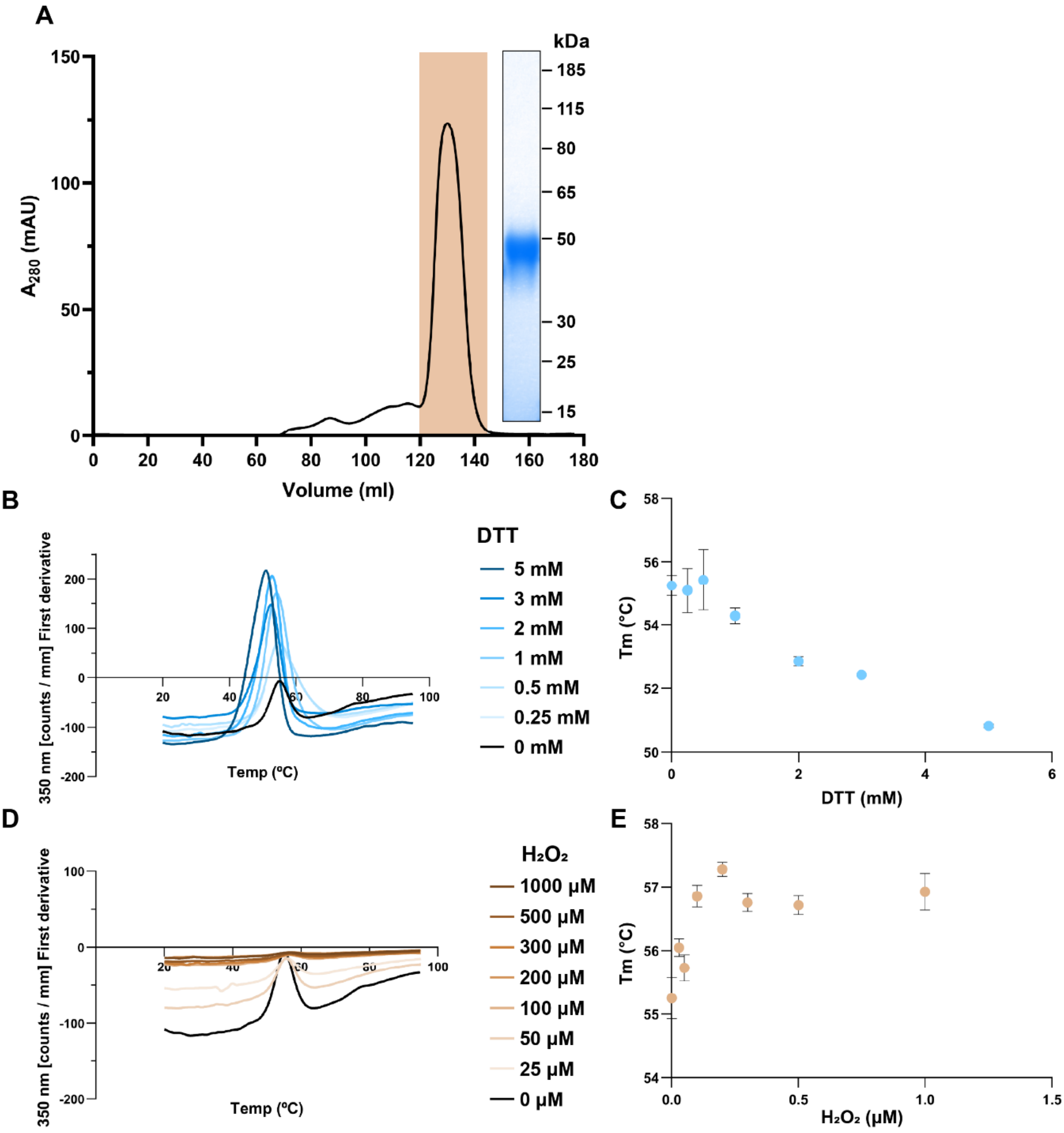


**Supplementary Figure 3**. **Thermal unfolding assay of CRK17 expressed in TniH5 cells upon treatment with increasing concentrations of DTT and $H_2O_2$**. A) After affinity chromatography, the His-tagged protein was separated by size-exclusion chromatography (SEC) on preparative-grade Superdex S200. CRK17 ECD eluted from 121-143.5 mL (orange peak, chromatogram, left). The SDS-PAGE gel (right) shows the final pooled and concentrated CRK17 ECD sample. Fluorescence was measured by nano differential scanning fluorimetry (DSF) using Prometheus Panta (NanoTemper Technologies GmbH). B, D) Thermal unfolding curves show 350 nm fluorescence emission of CRK17 at increasing temperature (0.5 °C/min). B, D) First derivative of thermal unfolding curves show 350 nm fluorescence emission of CRK17 at increasing temperature (0.5 °C/min). The inflexion point was used to determine the melting temperature ($T_m$). C, E) The $T_m$ as a function of increasing DTT and $H_2O_2$ concentrations. 0.4 mg/ml protein samples were prepared in pH 7.5 TBS supplemented with 1 mM $MgCl_2$ and 1 mM pluronic acid, with appropriate treatments (no treatment, DTT or $H_2O_2$).

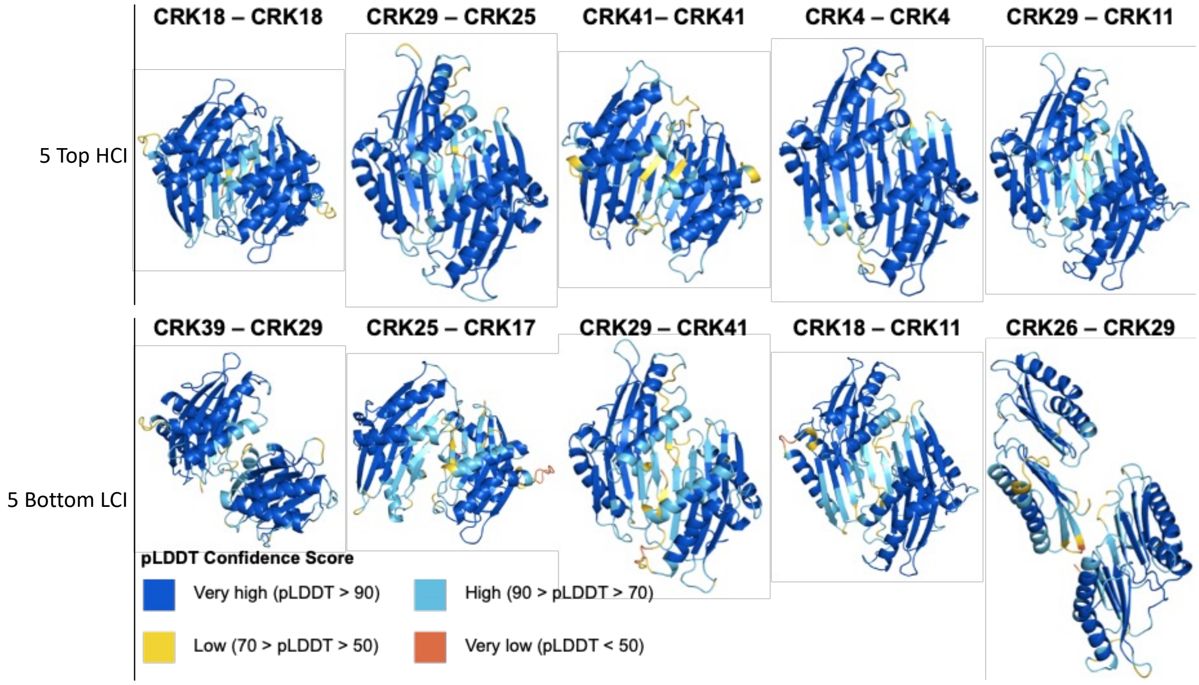


**Supplementary Figure 4. High- and Low-Confidence Interactions from No ROS conditions were run through AlphaFold Multimer v2.2.2.** Residues are coloured by predicted local distance difference test (pLDDT) scores.

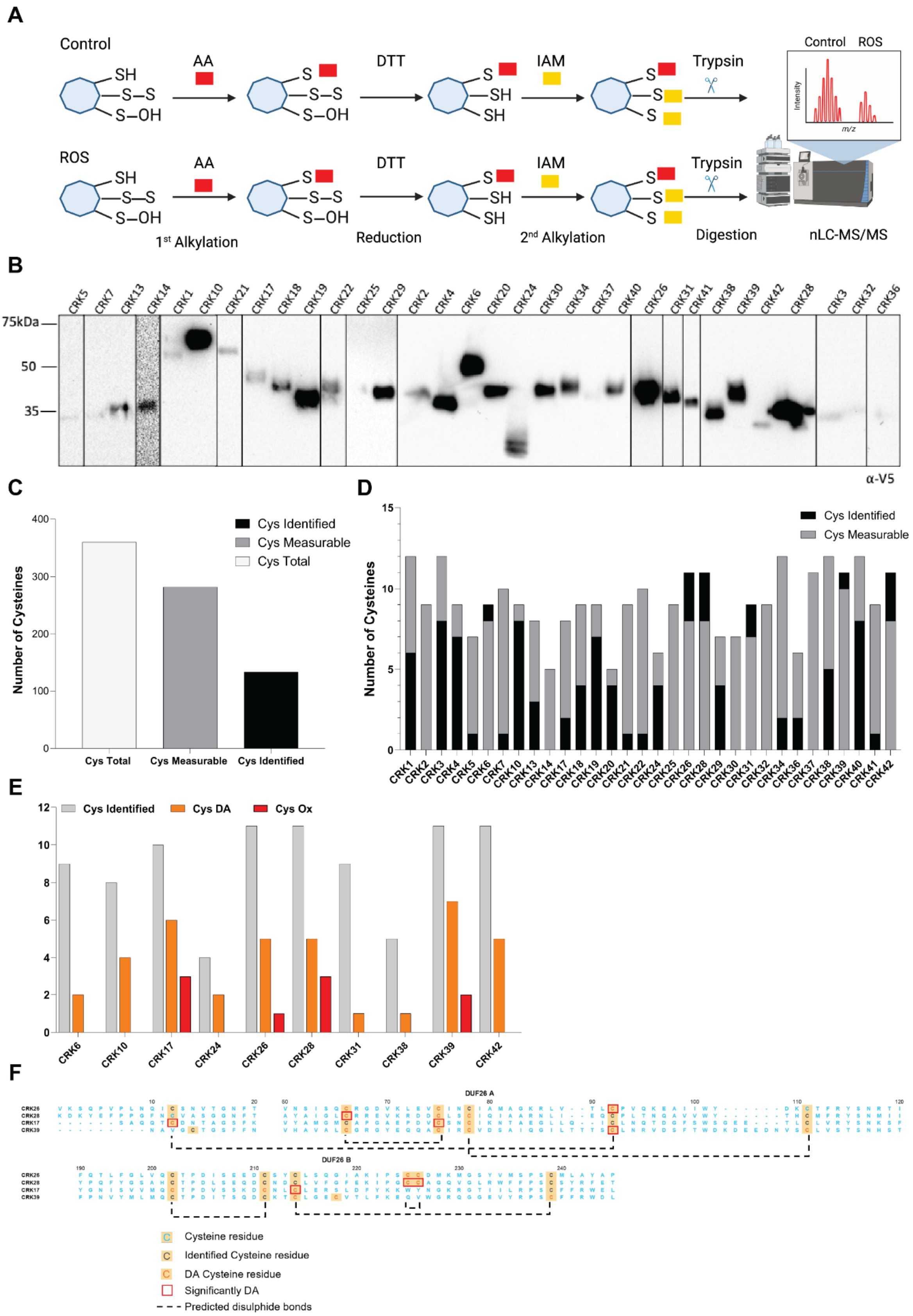


**Supplementary Figure 5. Identification of cysteine oxidative modifications.** A) Schematic overview of the differential alkylation method (DAM), which is a technique that labels reduced thiols using an alkylation agent, such as acrylamide (AA) or iodoacetamide (IAA). This is followed by the reduction (DTT) of all reversibly oxidised sites, followed by a second alkylation with another alkylating agent, such as iodoacetamide (IAM). This approach facilitates the identification and quantification of cysteine oxidation. B) Western blots of CRK ECDs expressed with minimal V5 tag and detected with anti-V5-HRP antibodies. C) Number of Total, Measurable and Identified cysteines for all tested CRK ECDs. D) Identified and Measurable Cysteines per individual tested CRK ECD. E) Number of identified, differentially labelled and oxidatively modified cysteines per CRK for which differential alkylation was measured. F) Amino acid alignment of CRK26, 28, 17 and 39, with indication of which cysteines we identified as DA and more oxidized upon $H_2O_2$ treatment.

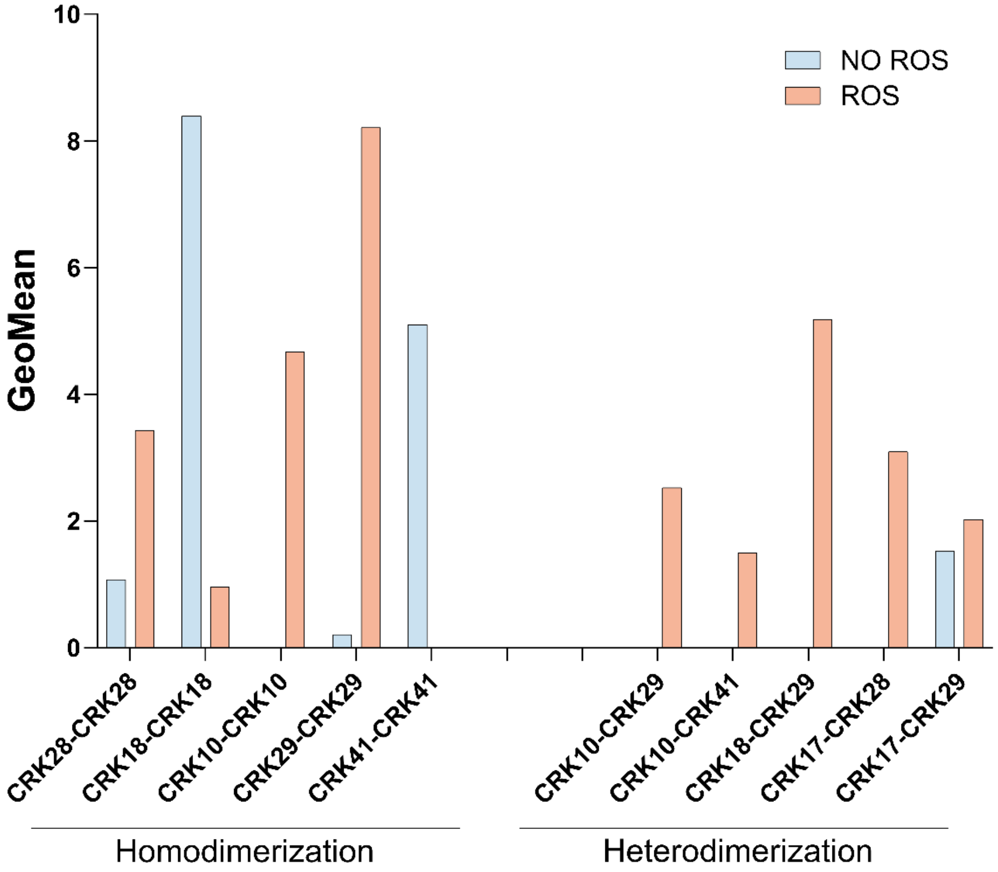


**Supplementary Figure 6. Interaction pairs from RIA$^{CRK}$ selected as candidates for in planta studies.** A) The top HCI pairs from RIA$^{CRK}$ were filtered by the highest GeoMean score, whether both interacting partners were expressed in senescent tissues and whether the interaction was modulated by ROS. The bar plots represent the Geomean score for both homodimeric interactions in the left and heterodimeric interactions in the right. In blue and orange, GeoMean in No ROS and ROS conditions respectively.

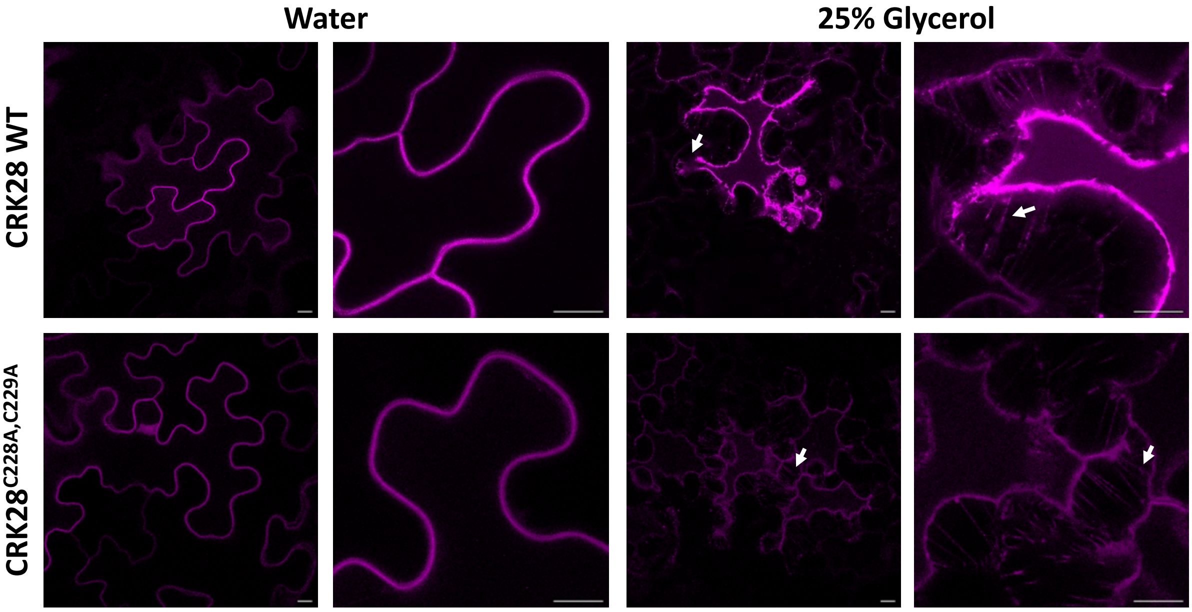


**Supplementary Figure 7. CRK28 and CRK28$^{C228A,C229A}$ localise to the plasma membrane.** Transient expression of 35S:CRK28$^{KD}$-3xFLAG-mScI and 35S:CRK28$^{C228A,C229A-KD}$-3xFLAG-mScI in *N. benthamiana* epidermal cells. After treatment with 25% glycerol, Hechtian strands are observed (white arrows) caused by detachment of the membrane from the cell wall upon plasmolysis, showing that CRK28 and CRK28$^{C228A,C229A}$ localise to the plasma membrane when transiently expressed in *N. benthamiana*. Scale bars are 10 μM.

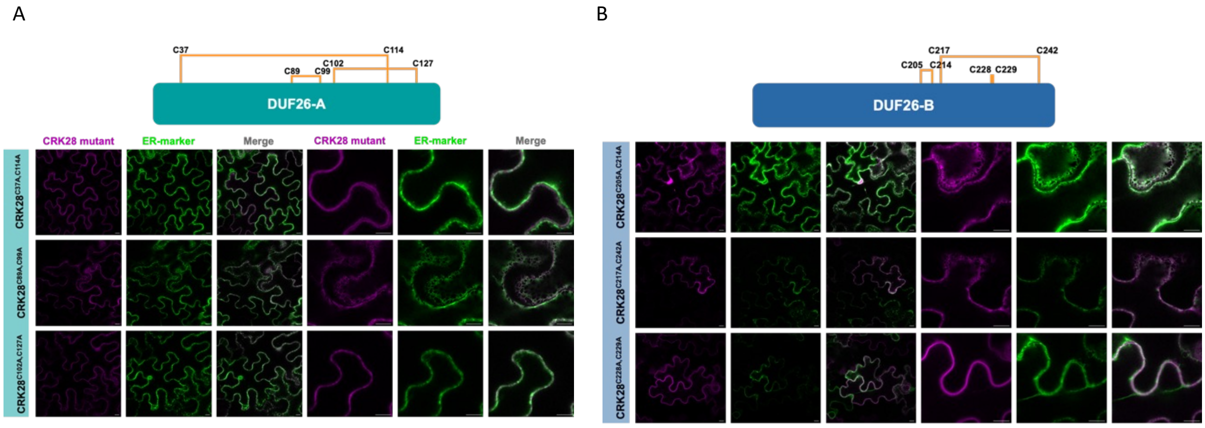


**Supplementary Figure 8. CRK28 cysteine residues are required for plasma membrane localisation.** Schematic of the disulphide bonds in CRK28-ECD DUF26-A and localisation of the CRK28 cysteine mutants, each of which lacks a disulphide bond. B) Schematic of the disulphide bonds in CRK28-ECD DUF26-B and localisation of the CRK28 cysteine mutants, each of which lacks a disulphide bond. Localisation to the ER is apparent for CRK28-ECD$^{C37A,C114A}$,CRK28-ECD$^{C89A,C99A}$, CRK28-ECD$^{C102A,C127A}$, CRK28-ECD$^{C205A,CRK214A}$ and CRK28-ECD$^{C217A,C242A}$. CRK28-ECD$^{C228A,C229A}$ shows uniform expression at the plasma membrane. CRK28 and all mutant variants have the K377N KD mutation. Scale bar is 10 μM.

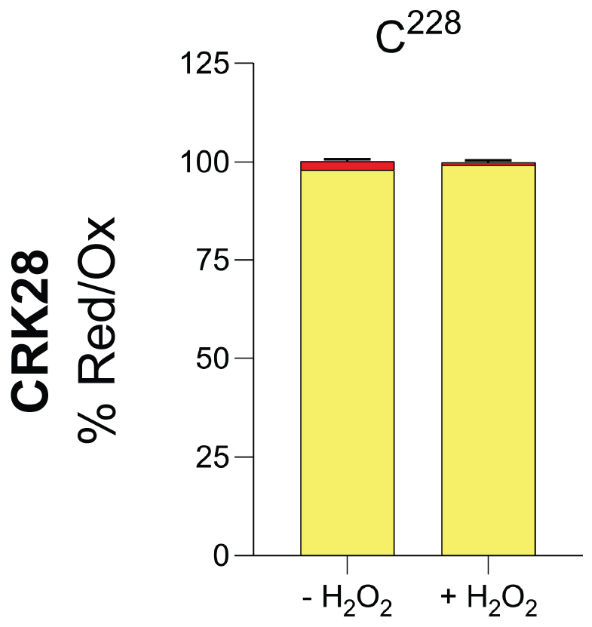


**Supplementary Figure 9. Cysteine 228 undergoes oxidative modification *in vivo*.** 35S:CRK28$^{KD}$-3xFLAG-mScI was transiently expressed in *Nicotiana bethamiana*, treated with 4 mM $H_2O_2$ or $MgCl_2$ (mock). CRK28 was enriched using anti-FLAG beads, and samples were subjected to DAM followed by MS to measure cysteine oxidation. In yellow, the percentage of oxidized forms of cysteine (S-S/-SOH), in red, the percentage of reduced cysteine (-SH).

A

| Cysteine | RSA (%) | Ox *in vitro* | Ox *in vivo* | Cysteine mutant | Localization |
|---|---|---|---|---|---|
| C37 | 54 | | | C37A,C114A | ER/PM |
| C89 | 5 | Yes | | | |
| C99 | 1 | | | C89A, C99A | ER |
| C102 | 15 | | | | |
| C114 | 2 | | | C102A, C127A | ER |
| C127 | 3 | | | | |
| C205 | 0 | | | C205A, C214A | ER |
| C214 | 0 | | | | |
| C217 | 0 | | | C217A, C242A | ER |
| C228 | 47 | Yes | Yes | | |
| C229 | 0 | Yes | | C228A, C229A | PM |
| C242 | 5 | | | | |

B

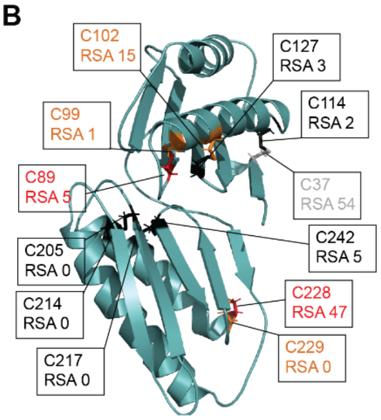


**Supplementary Figure 10. The summary table presents the effects of disulphide bonds on the stability and localisation of CRK28 and indicates which cysteines are oxidatively modified.** For each cysteine residue in the CRK28-ECD AlphaFold model, solvent accessibility was calculated as a percentage of the available surface area. The presence of oxidative modification, *in vitro* or *in vivo*, is indicated. Localisation of full-length CRK28 mutant variants in *N. benthamiana* is indicated in the table. The positions of the disulphide bonds in the AlphaFold structure of CRK28-ECDs are marked in grey.

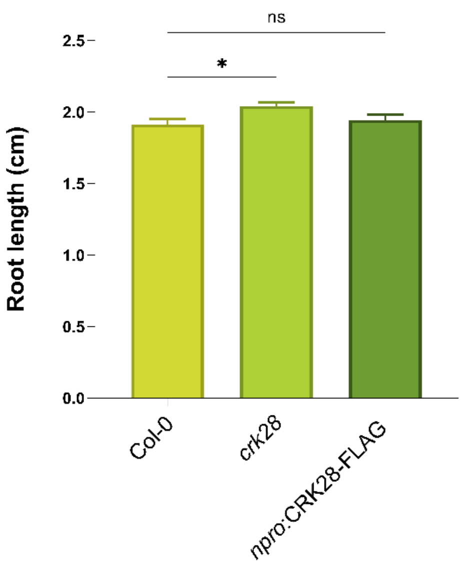


**Supplementary Figure 11. CRK28 does not affect early plant development.** Root growth measurements of *crk28* and *npro*:CRK28-FLAG plants grown for 14 days under optimal conditions (long day, 16 hours of light and 8 hours of darkness, 21 °C). Asterisks denote statistical significance in one-way ANOVA of each sample compared with Col-0. ns = not significant, * p-value < 0.5, **** p-value < 0.0001.

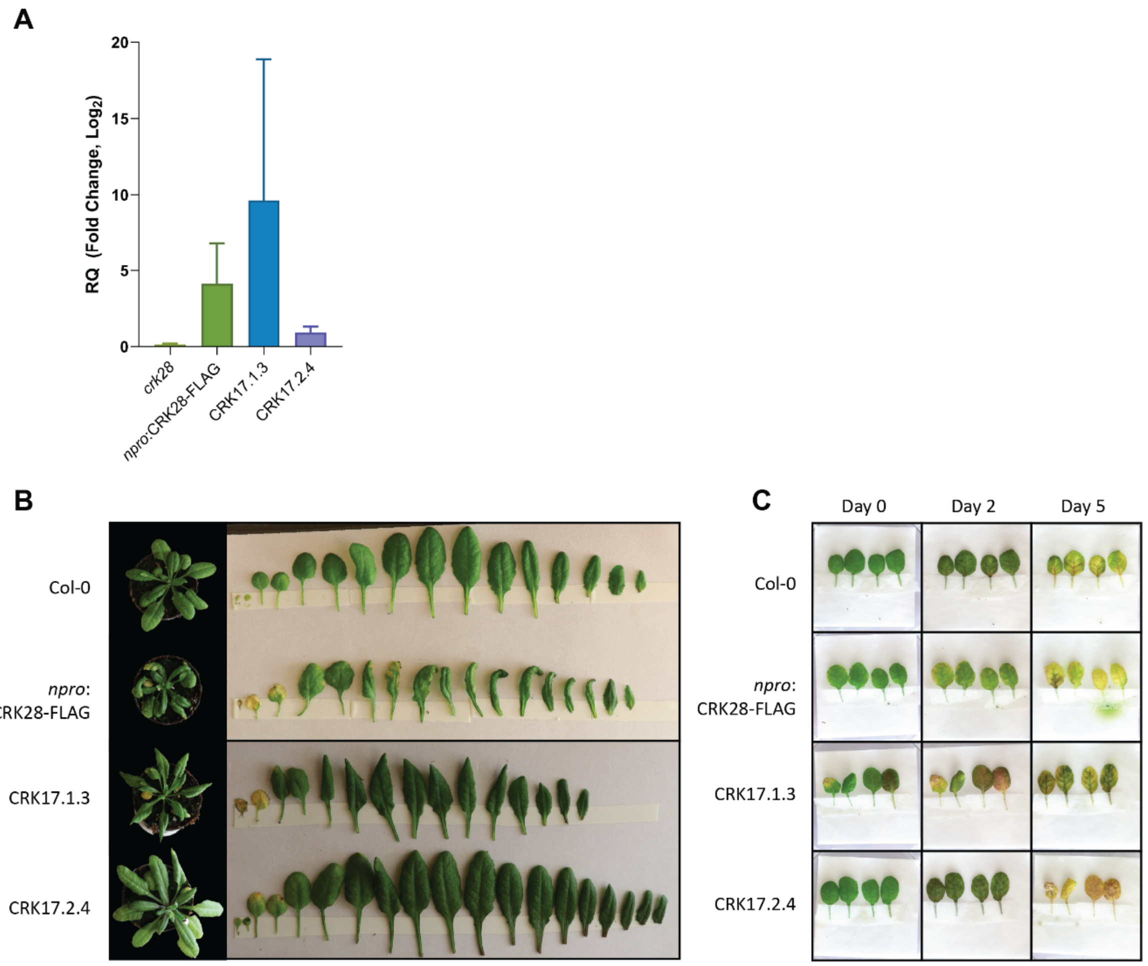


**Supplementary figure 12. Overexpression of CRK17 resembles the *npro*:CRK28-FLAG morphology.** A) Expression of CRK28 in *crk28* T-DNA mutant and *npro*:CRK28-FLAG lines, and CRK17 in 35S: CRK17 lines, compared to Col-0. Expression was measured by qRT-PCR as described in Materials & Methods. Expression was measured in at least 3 independent leaves per sample. B) 5-week-old Arabidopsis rosettes and leaf panels of representative Col-0, *npro*:CRK28-FLAG and 35S:CRK17 lines. C) Leaf detachment assay on the 5th and 6th leaves of 5-week-old plants. Pictures taken on Day 0 and Day 2 to assess senescence progression. Representative images of samples from the 5th leaf (left) and 6th leaf (right) in each square.

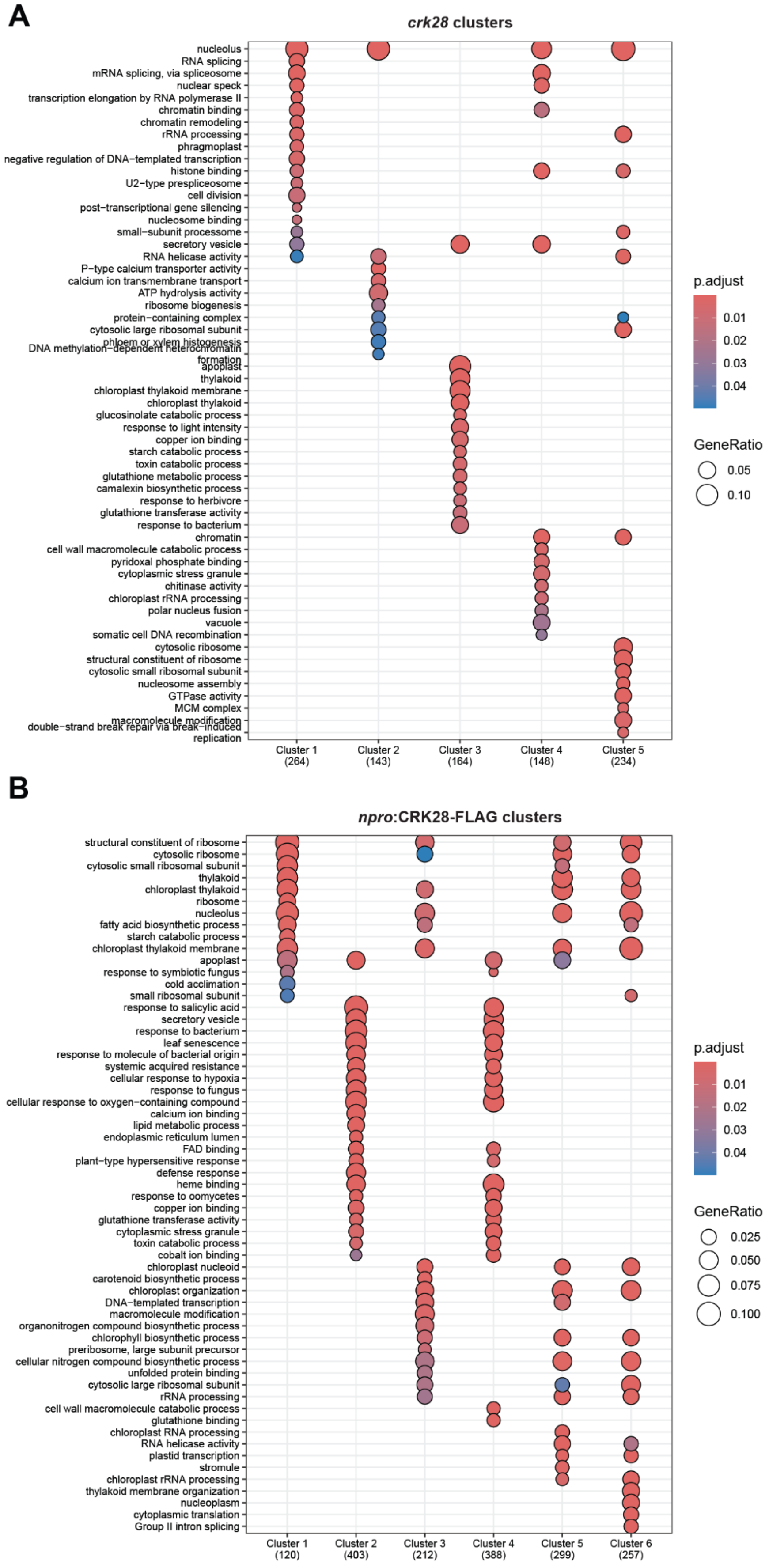
A
crk28 clusters
nucleolus
RNA splicing
mRNA splicing, via spliceosome
nuclear speck
transcription elongation by RNA polymerase II
chromatin binding
chromatin remodeling
rRNA processing
phragmoplast
negative regulation of DNA−templated transcription
histone binding
U2−type prespliceosome
cell division
post−transcriptional gene silencing
nucleosome binding
small−subunit processome
secretory vesicle
RNA helicase activity
P−type calcium transporter activity
calcium ion transmembrane transport
ATP hydrolysis activity
ribosome biogenesis
protein−containing complex
cytosolic large ribosomal subunit
phloem or xylem histogenesis
DNA methylation−dependent heterochromatin formation
apoplast
thylakoid
chloroplast thylakoid membrane
chloroplast thylakoid
glucosinolate catabolic process
response to light intensity
copper ion binding
starch catabolic process
toxin catabolic process
glutathione metabolic process
camalexin biosynthetic process
response to herbivore
glutathione transferase activity
response to bacterium
chromatin
cell wall macromolecule catabolic process
pyridoxal phosphate binding
cytoplasmic stress granule
chitinase activity
chloroplast rRNA processing
polar nucleus fusion
vacuole
somatic cell DNA recombination
cytosolic ribosome
structural constituent of ribosome
cytosolic small ribosomal subunit
nucleosome assembly
GTPase activity
MCM complex
macromolecule modification
double−strand break repair via break−induced replication
Cluster 1 (264)
Cluster 2 (143)
Cluster 3 (164)
Cluster 4 (148)
Cluster 5 (234)
p.adjust
0.01
0.02
0.03
0.04
GeneRatio
0.05
0.10
B
npro:CRK28-FLAG clusters
structural constituent of ribosome
cytosolic ribosome
cytosolic small ribosomal subunit
thylakoid
chloroplast thylakoid
ribosome
nucleolus
fatty acid biosynthetic process
starch catabolic process
chloroplast thylakoid membrane
apoplast
response to symbiotic fungus
cold acclimation
small ribosomal subunit
response to salicylic acid
secretory vesicle
response to bacterium
leaf senescence
response to molecule of bacterial origin
systemic acquired resistance
cellular response to hypoxia
response to fungus
cellular response to oxygen−containing compound
calcium ion binding
lipid metabolic process
endoplasmic reticulum lumen
FAD binding
plant−type hypersensitive response
defense response
heme binding
response to oomycetes
copper ion binding
glutathione transferase activity
cytoplasmic stress granule
toxin catabolic process
cobalt ion binding
chloroplast nucleoid
carotenoid biosynthetic process
chloroplast organization
DNA−templated transcription
macromolecule modification
organonitrogen compound biosynthetic process
chlorophyll biosynthetic process
preribosome, large subunit precursor
cellular nitrogen compound biosynthetic process
unfolded protein binding
cytosolic large ribosomal subunit
rRNA processing
cell wall macromolecule catabolic process
glutathione binding
chloroplast RNA processing
RNA helicase activity
plastid transcription
stromule
chloroplast rRNA processing
thylakoid membrane organization
nucleoplasm
cytoplasmic translation
Group II intron splicing
Cluster 1 (120)
Cluster 2 (403)
Cluster 3 (212)
Cluster 4 (388)
Cluster 5 (299)
Cluster 6 (257)
p.adjust
0.01
0.02
0.03
0.04
GeneRatio
0.025
0.050
0.075
0.100

**Supplementary figure 13.** GO-term analysis was performed for protein hit clusters in Fig. 4 C for A) *crk28* and B) *npro*:CRK28-FLAG plants.

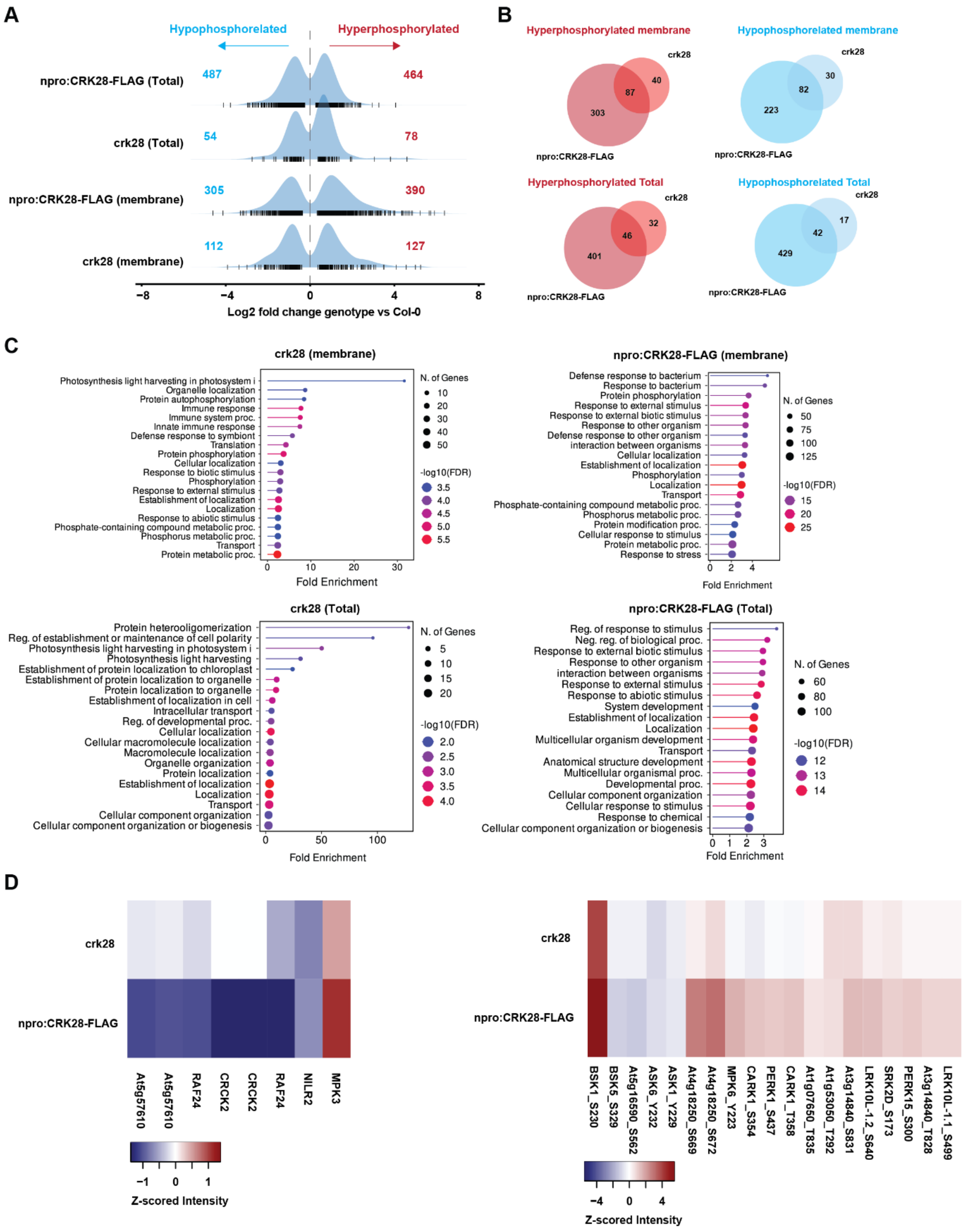


**Supplementary Figure 14. CRK28 rewires the phosphoproteome in 5-week-old plants.** A) Distribution histograms of significantly differential phosphosites (FDR ≤ 0.05) comparing genotype vs Col-0 across the total and membrane enriched fractions. Numbers of hyper- or hypo-phosphorylated sites are indicated. B) Venn diagrams showing the numbers of significant hyper- and hypo-phosphorylated peptides identified in the total proteome or membrane-enriched fractions of *crk28* and npro:CRK28-FLAG. C) Gene Ontology (GO) enrichment of significant hyper- and hypo-phosphorylated peptides identified in the total proteome or membrane-enriched fractions of *crk28* and npro:CRK28-FLAG. D) Hyper- and hypo-phosphorylation of the kinases identified in the total and membrane-enriched proteomes of *crk28* and npro:CRK28-FLAG. The full lists are available in the Supp. Tables 12-17.
